# Local chromatin decompaction shapes mitotic chromosome landscape

**DOI:** 10.1101/2024.06.30.601459

**Authors:** Sanki Tashiro, Hideki Tanizawa, Ken-ichi Noma

## Abstract

Unregulated activation of transcription on mitotic chromosomal arms can be detrimental to faithful chromosome segregation^1–3^, although a subset of genes are up-regulated during mitosis^4^, raising the question as to how gene activation is coordinated in the context of mitotic chromosome reorganization. Here we investigate the fine structure of mitotic chromosomes in fission yeast *Schizosaccharomyces pombe* and reveal that intense decompaction of local chromatin upon mitotic gene activation helps shape the mitotic chromosome landscape. More specifically, we show that binding of mitosis-specific transcription factors, Ace2 and MluI binding factor (MBF)^5,6^, to gene promoters followed by transcriptional elongation can locally induce chromatin decompaction during mitosis. Interestingly, local decompaction-competent genes can establish chromatin boundaries that demarcate long-range contact domains, indicating tight coupling between local and global genome organizations. Furthermore, efficient local decompaction requires enough transcriptional elongation distance and histone removal, providing a mechanistic insight into the decompaction-associated domain boundary formation. Given that the demarcation of large, self-associating domains is critical for faithful chromosome segregation^7^, we propose that decompaction-competent genes can be a key determinant of mitotic chromosome configuration, thereby underpinning the maintenance of genome integrity over generations, and thus their distribution along chromosomes can be subject to evolutionary selection.

## INTRODUCTION

During mitosis, chromosomal DNA in eukaryotic cells is subject to intense condensation, which confers duplicated sister chromatids mechanical stiffness necessary for their accurate segregation into daughter cells^8^. Central to this process is highly conserved condensin, a pentameric protein complex that contains structural maintenance of chromosomes (SMC) proteins. Based on high-throughput chromosome conformation capture (Hi-C) combined with polymer simulations, it has been proposed that condensin can organize miotic chromosome structures such that chromosomal DNA is folded into helically arranged scaffolds with protruded loops ^9^, which agrees with previous microscopic observation of the mitotic chromosome structure^10,11^. However, electron tomography of mitotic chromosomes detected no such regular arrangement of chromatin fibers^12–14^, challenging the view of highly ordered chromosome arrangements. Thus, the basic mechanism of how long genomic DNA is assembled into compact mitotic chromosomes remains obscure.

Along with the structural reorganization of chromosomes, transcription in mitotic cells is generally shut down by the clearance of RNA polymerases from chromatin^1^. Failure in this process can lead to deleterious outcomes such as lagging chromosomes and chromosome bridges^1–3^. Even though mitotic transcriptional repression has been well described in many organisms, a sensitive assay using a cell-permeable nucleotide analog has unveiled that transcription at many genes remains active, albeit at low levels, during mitosis^4^. Interestingly, this study also showed that a subset of genes are even mitotically up-regulated, demonstrating that mitotic chromosomes are not transcriptionally silent entities. It remains unclear, however, how local chromatin structure at mitotically activated genes is coordinated in the context of extensive chromosomal reorganization during mitosis.

In fission yeast mitosis, as in higher eukaryotes, condensin plays an essential role in chromosome condensation and segregation^15,16^, while a subset of genes are up-regulated^5,6^. Interestingly, even though global repression of transcription during mitosis is not evident in this organism, transcription inhibition can promote chromosomal segregation fidelity by suppressing a partial loss of condensin function^17,18^. Thus, fission yeast can serve as an ideal model organism to study how active gene transcription is orchestrated with the mitotic chromosome assembly. Using fission yeast, we previously analyzed time-dependent alterations of the mitotic chromosome structure and described the dynamic assembly and disassembly of condensin-dependent, large self-associating chromatin domains ranging from 300 kb to 1 Mb in size^19^. Our previous study also shows that condensin can be recruited to some gene loci by its physical interaction with a specific transcription factor, at which boundaries of condensin-mediated large domains that are necessary for faithful chromosomal segregation are formed^7^. In the present study, we turn to fine structures of local, gene-sized chromatin along mitotic chromosomes by modifying the existing Hi-C protocol so that short-range genomic DNA contacts can be captured. As a result, we find unusually intense decompaction of local chromatin at a limited number of mitotically activated genes. Interestingly, this local decompaction is tightly connected to the formation of chromosome-sized domain boundaries, suggesting a possible role of mitotically activated genes in the arrangement of mitotic chromosome configuration.

## RESULTS

### Local hyper-decompaction at limited gene loci during mitosis

In order to map local DNA contacts of mitotic chromosomes at high resolution, we made simple modifications in the conventional in situ Hi-C method (**Extended Data Fig. 1**, see also **Supplementary Note**). Similar to previous studies^20,21^, we used multiple restriction enzymes, MboI, HinfI, and MluCI, so that chromosomal DNA was digested into much smaller fragments compared to single MboI enzyme treatment (**Extended Data Fig. 1a,b**). In addition, we explored chromatin fixation conditions as in the previous study^22^ and found that double fixation combining paraformaldehyde (pFA) and disuccinimidyl glutarate (DSG) resulted in better detection of short-range contacts (**Extended Data Fig. 1c-f**). This double fixation, triple digestion condition was also successful in the detection of long-range contact with marginally less efficacy, being able to reproduce cohesin- and condensin-mediated assembly of self-associating domains^19,23^ (**Extended Data Fig. 1g**).

By applying this modified Hi-C approach, we sought to determine the fine structures of mitotic chromosomes in fission yeast. To prepare mitotically synchronized cell cultures, we utilized the *cdc25-22* temperature-sensitive mutant allele, which allows cell cycle arrest at the G2/M phase transition at a restrictive temperature followed by synchronous release into mitosis at a permissive temperature. After verifying the synchronous cell cycle progression by monitoring mitotic spindles and cell septation, which are indicative of M and S phases, respectively (**Extended Data Fig. 2a**), we performed Hi-C every 10 minutes from 20 to 70 minutes after the release. Consistent with a previous finding^19^, the modified Hi-C protocol detected the dynamic assembly and disassembly of the condensin-mediated large domains (300 kb to 1 Mb in size) during mitosis (from 20 to 50 minutes) and after mitosis (from 50 to 70 minutes), respectively (**Fig. 1a** and **Extended Data Fig. 2b,c**). In addition, we observed a global decline in short-range contacts during mitosis, as evidenced by the decreased contact scores along the diagonal in difference Hi-C matrices from 20 to 50 minutes (**Fig. 1b**), which is at least partly reflective of the loss of cohesin-mediated domains^7,19,23^. These results suggest that chromosomal DNA during mitosis is compacted in terms of long-range contacts but concomitantly decompacted in the local context. Interestingly, we noticed that the degree of decompaction was far from uniform across the genome; an extraordinary decrease in very local contacts was observed only at tens of gene loci across the genome, as exemplified by the *eng1* and *SPAC343.20* gene loci (**Fig. 1c**), demonstrating that a limited number of gene loci are subject to hyper-decompaction amid the global decline in short-range DNA contacts during mitosis.

**Fig. 1.**
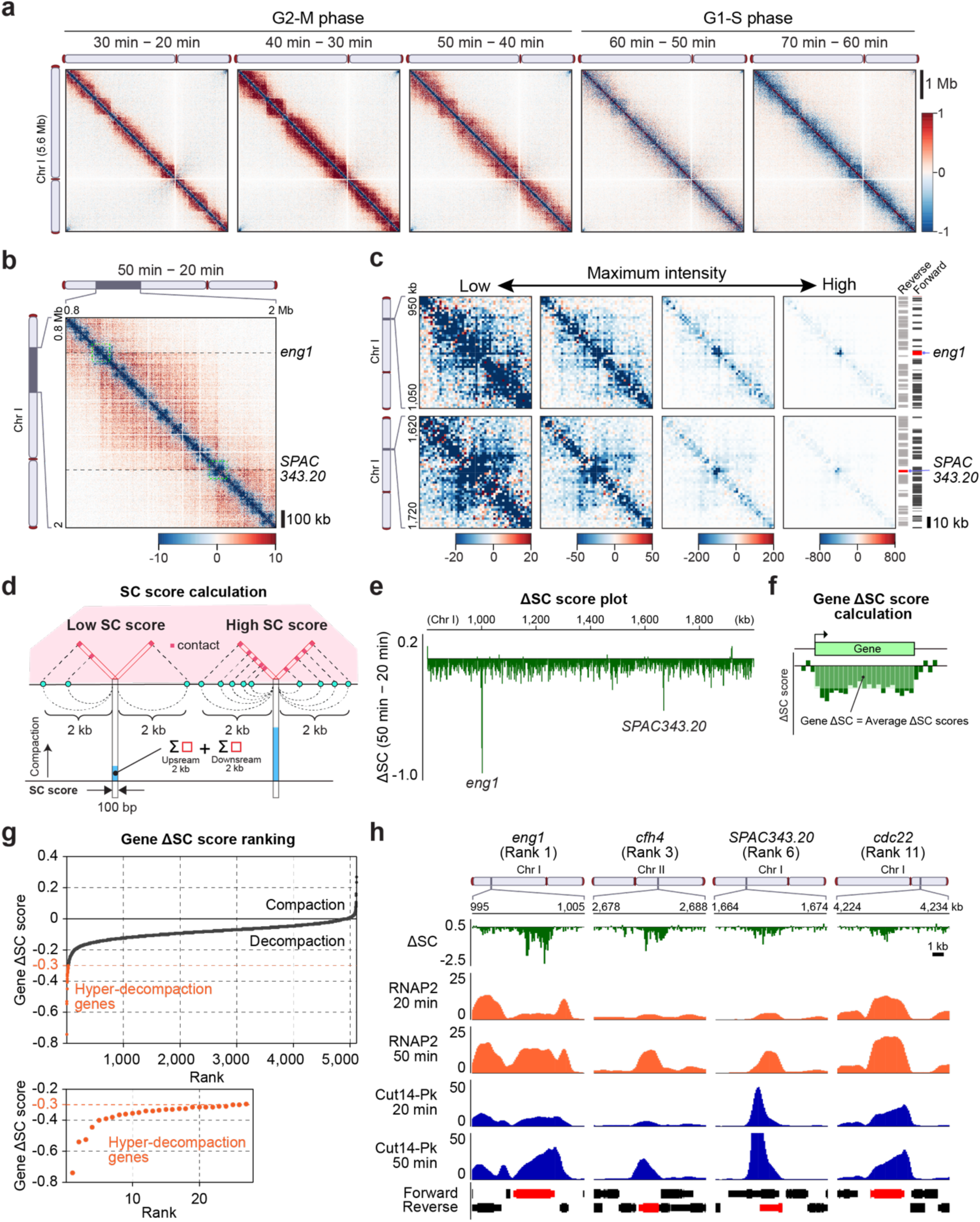
Intense local decompaction at limited gene loci during mitosis. **a**, Time-dependent alteration in genomic contacts during mitosis. Mitotically synchronized *cdc25-22* cells were fixed at 20, 30, 40, 50, 60, and 70 minutes after release from the G2/M transition, and in situ Hi-C experiments were performed. Differential contacts between two neighboring time points across chromosome I are shown. The increased and decreased contact scores over time are indicated by red and blue colors, respectively. For more information on cell cycle progression after the *cdc25-22* block release, see **Extended Data Fig. 2a**. **b**, Differential contacts between 20 and 50 minutes at a 1.2-Mb region of chromosome I that contains the *eng1* and *SPAC343.20* genes. The difference map (green dotted boxes) is enlarged in Fig. 1c. **c**, The same Hi-C difference maps as Fig. 1b but with the *eng1* or *SPAC343.20* gene loci enlarged. Very few blue signals survive even at color scales with an unusually high maximum intensity. On the right side of the Hi-C difference maps, the positions of the *eng1* and *SPAC343.20* genes are indicated by the red boxes, and all the other genes located in these regions are shown as black and gray boxes. **d**, Calculation schemes of short-range compaction (SC) scores and differential SC scores between 20 and 50 minutes (ΔSC). The total score of contacts within 2 kb from each 100-bp bin is defined as an SC score at that bin. **e**, The ΔSC score plot at the same 1.2-Mb region as Fig. 1b, which contains the *eng1* and *SPAC343.20* genes. **f**, A schematic of the gene ΔSC score definition. The average ΔSC score of 100-bp bins that overlap a transcribed region, including the 5′- and 3′-untranslated regions, in addition to the coding sequence of each protein-coding gene, is defined as the gene ΔSC score. **g**, Gene ΔSC score ranking and hyper-decompaction genes. In the upper panel, all the protein-coding genes were ranked based on the gene ΔSC scores and plotted in ascending order so that highly decompacted genes were to be top-ranked. The hyper-decompaction genes (a gene ΔSC score less than -0.300) are indicated by dots in orange, and their ΔSC score rank plot is shown in the lower panel. **h**, ΔSC score, RNA polymerase II (RNAP2) ChIP-seq, and Cut14-Pk ChIP-seq plots at example hyper-decompaction gene loci. The gene ΔSC score ranks are indicated above the plots.

To systematically identify gene loci that undergo hyper-decompaction during mitosis, we defined a short-range compaction (SC) score, which counts total contacts within 2 kb using 100-bp bin size (**Fig. 1d**). The differential SC scores between 20 and 50 minutes, referred to as ΔSC scores, were determined to detect the alteration in local compaction levels upon mitosis (**Fig. 1d**). As observed in the Hi-C maps in **Fig. 1c**, the unusually intense decrease in SC scores was detected at the *eng1* and *SPAC343.20* genes by calculating the ΔSC scores (**Fig. 1e**). Moreover, we determined ΔSC scores for each protein-coding gene, as in **Fig. 1f**, and ranked all the genes according to the gene ΔSC scores in ascending order so that intensely decompacted genes were to be top-ranked (**Fig. 1g**). Strikingly, there is a sharp drop in ΔSC scores in top-ranked genes in the ranking plot, and we defined the top 0.5% genes with the lowest gene ΔSC scores as hyper-decompaction genes in this study (27 genes with a gene ΔSC score less than -0.300, see also **Supplementary Table 1**). Notably, local decompaction occurred predominantly within transcribed regions but not at intergenic sequences (**Fig. 1h** and **Extended Data Fig. 3a**). Moreover, these hyper-decompaction genes were mitotically up-regulated genes associated with condensin at their 3′ ends in many cases, which was verified by chromatin immunoprecipitation followed by deep sequencing (ChIP-seq) (**Fig. 1h** and **Extended Data Fig. 3b**), suggesting that mitotic gene up-regulation relates to hyper-decompaction.

### Ace2-driven transcription can induce local decompaction and long-range domain boundaries

Proper progression of mitosis in fission yeast cells requires timely expression of several transcription factors at different mitotic stages^5,6^. Interestingly, one such transcription factor, Ace2, seems to target many of the local hyper-decompaction genes identified here, such as *eng1*, *cfh4*, and so forth^24^. To substantiate this observation, we performed ChIP-seq for Ace2 in mitotically synchronized cells and found that Ace2 was indeed located at promoters of many hyper-decompaction genes (**Fig. 2a,b**). This finding urged us to investigate the possible role of the Ace2 transcription factor in regulating local compaction. Because Ace2 depletion likely affects cell cycle progression (Rustici et al. 2004), we instead introduced small deletions of the two Ace2-binding motifs (5′-CCAGCC-3′ x2) in the promoter region of the *eng1* gene (*Peng1*)^25^ to compromise the local Ace2 binding. In addition to this, we also constructed cells carrying an 11-bp deletion of the putative TATA box (5′-AATAATATAAA-3′) in *Peng1* to test the effect of reduced transcription activity. Expectedly, our reverse transcription followed by quantitative polymerase chain reaction (RT-qPCR) showed that these mutations, referred to as *eng1-motif*ΔΔ and *eng1-TATA*Δ, largely compromised *eng1* mRNA expression during mitosis (**Extended Data Fig. 4a**), even though the Ace2 binding at *Peng1* was only mildly affected in *eng1-TATA*Δ (**Extended Data Fig. 4b**). We performed Hi-C using mitotically synchronized cells at 20 and 50 minutes after G2/M block release and found that the intense decompaction observed at the *eng1* gene in the wild type was diminished in both *eng1* mutants (**Fig. 2c**), suggesting that binding of the Ace2 transcription factor at a promoter followed by transcription up-regulation is necessary for efficient local decompaction at *eng1* during mitosis.

**Fig. 2.**
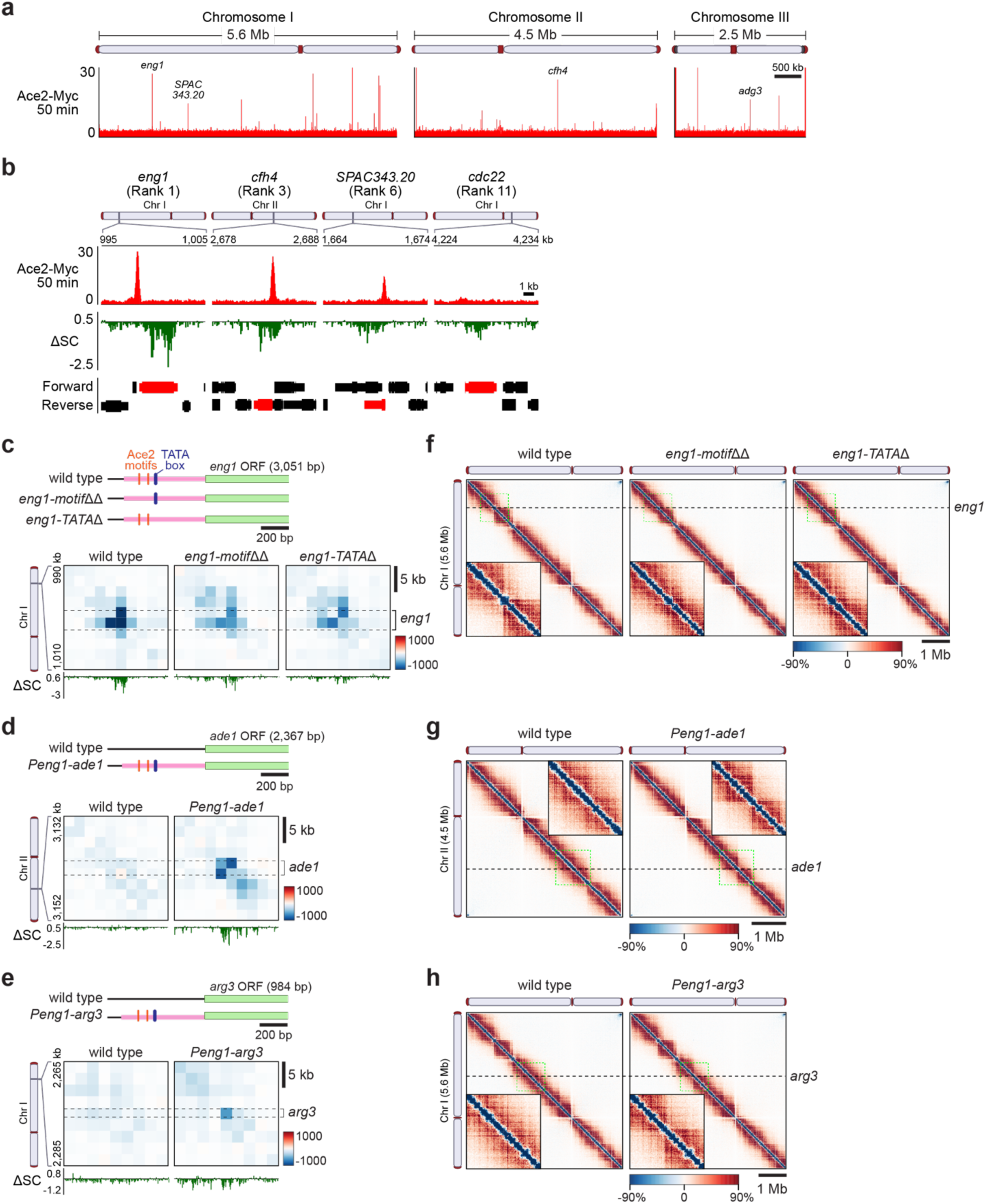
Ace2-driven *eng1* promoter can induce local decompaction and boundary formation of mitotic chromatin domains. **a**, A genome-wide binding profile of Ace2 transcription factor during mitosis. Mitotically synchronized *cdc25-22* cells expressing Ace2-Myc from the endogenous promoter were fixed at 50 minutes after G2/M block release, and ChIP-seq was performed using anti-Myc antibody. **b**, Ace2-Myc ChIP-seq plots at the example hyper-decompaction genes as in Fig. 1h. The same ΔSC score plots as in Fig. 1h are again shown for comparison. **c**, Defective decompaction at *eng1* upon deletion of the two Ace2 binding motifs (*eng1-motif*ΔΔ) or the TATA box (*eng1-TATA*Δ). The upper panel illustrates the *eng1* gene locus in the wild type and *eng1* mutants. The positions of the *eng1* promoter, Ace2 binding motifs, and putative TATA box are indicated by pale red, orange, and blue boxes, respectively. The lower panel shows Hi-C difference maps between the wild type and the mutants. Mitotically synchronized *cdc25-22* cells with the wild-type or mutant *eng1* gene were fixed at 20 and 50 minutes after release from the G2/M block, and Hi-C experiments were performed. After each Hi-C matrix was prepared, the differential contacts between 20 minutes (G2 phase-prophase) and 50 minutes (metaphase-anaphase) were calculated for each strain, and a 20-kb region of chromosome I that contains the *eng1* gene is displayed. Below each Hi-C difference map is its ΔSC score plot in the same region. **d**, De novo hyper-decompaction at ectopic sites upon insertion of the *eng1* promoter (*Peng1*) at the *ade1* (*Peng1-ade1*) gene. The upper panel illustrates the *ade1* gene locus in the wild-type and *Peng1* insertion strains. The differential contacts between 20 and 50 minutes, as in Fig. 2c but at a 20-kb region of chromosome II, are displayed, and below each Hi-C difference map is its ΔSC score plot in the same region. **e**, The same analysis as in Fig. 2d but with the *arg3* gene as an ectopic *Peng1* insertion site (*Peng1-arg3*). **f**, Defective boundary formation at the *eng1* locus in the *eng1-motif*ΔΔ and *eng1-TATA*Δ mutants. The Hi-C difference maps, as in Fig. 2c, are shown for the whole chromosome I. An enlarged view of the 1-Mb region that contains the *eng1* gene, as indicated by a dotted line box in green, is shown as an inset. **g**, De novo boundary formation in the *Peng1-ade1* strain. The difference maps, as in Fig. 2d, are shown for the whole chromosome II. An enlarged view of the 1-Mb region that contains the *ade1* gene, as indicated by a dotted line box in green, is shown as an inset. **h**, The same analysis as in Fig. 2g but with the *Peng1-arg3* strain.

The necessity of the Ace2-driven transcription for local decompaction during mitosis led us to examine whether the *Peng1* sequence can actively decompact local chromatin when ectopically placed at unrelated genes. We picked constitutively active *ade1* and *arg3* genes and inserted *Peng1* at the immediate upstream of the coding sequence so that these genes were to be activated by Ace2. Indeed, the *Peng1* insertions forced these genes to undergo mitotic activation (**Extended Data Fig. 4c**). We also performed Hi-C and found that both *Peng1* insertions at the *ade1* and *arg3* genes locally induced a remarkable decrease in short-range DNA contacts at these genes during mitosis (**Fig. 2d,e**), suggesting that *Peng1* can decompact local chromatin during mitosis no matter what genes it regulates.

Strikingly, these small genetic alterations examined in **Fig. 2c-e** caused large-scale alterations in mitotic chromosome conformation. As shown in **Fig. 2f**, the *motif*ΔΔ and *TATA*Δ mutations at the endogenous *eng1* promoter caused enhanced contacts between the two adjoining condensin-mediated, self-associating domains located upstream and downstream of the *eng1* gene locus ^19^ (**Extended Data Fig. 2c)**. In other words, the boundary that is inhibitory to long-range DNA contacts between these two condensin-mediated domains was compromised in these mutants.

In addition, the *Peng1* insertions at *ade1* and *arg3* led to *de novo* formation of boundaries that suppressed long-range DNA contacts (**Fig. 2g,h**). Collectively, these results indicate that the Ace2-driven promoter *Peng1* can not just induce mitotic transcription and local chromatin decompaction but also inhibit long-range chromosomal DNA contacts between its flanking domains.

### Local decompaction activity is intimately connected to boundary formation

The above Hi-C results suggest that these two phenomena in short- and long-range DNA contacts are mechanistically connected. To explore this possibility, mutated *Peng1* that lacks the Ace2 binding motifs [*Peng1*(*motif*ΔΔ)] or the TATA box [*Peng1*(*TATA*Δ)] was placed at the *ade1* gene locus as in the *Peng1-ade1* strain analyzed in **Fig. 2d,g**. After verifying that the mitotic activation of the *ade1* gene observed in *Peng1-ade1* cells was abrogated in the mutant *Peng1-ade1* strains (**Extended Data Fig. 4d**), we performed Hi-C to examine whether short- and long-range contacts around the *ade1* gene locus were affected in these genetic backgrounds. In terms of local contacts, the intense decompaction observed at the *Peng1-ade1* locus was compromised in the *Peng1*(*motif*ΔΔ)*-ade1* strain and also affected in the *Peng1*(*TATA*Δ)*-ade1* strain to a lesser extent (**Fig. 3a**). On the other hand, when it comes to long-range contacts, the boundary-forming activity of the *Peng1* insertions was almost totally abolished in the *Peng1*(*motif*ΔΔ)*-ade1* strain and was weakened in the *Peng1*(*TATA*Δ)*-ade1* strain (**Fig. 3b**). This apparent correlation between short- and long-range DNA contact alterations upon insertion of the *Peng1* variants again indicates the intimate mechanistic connection between the two phenomena.

**Fig. 3.**
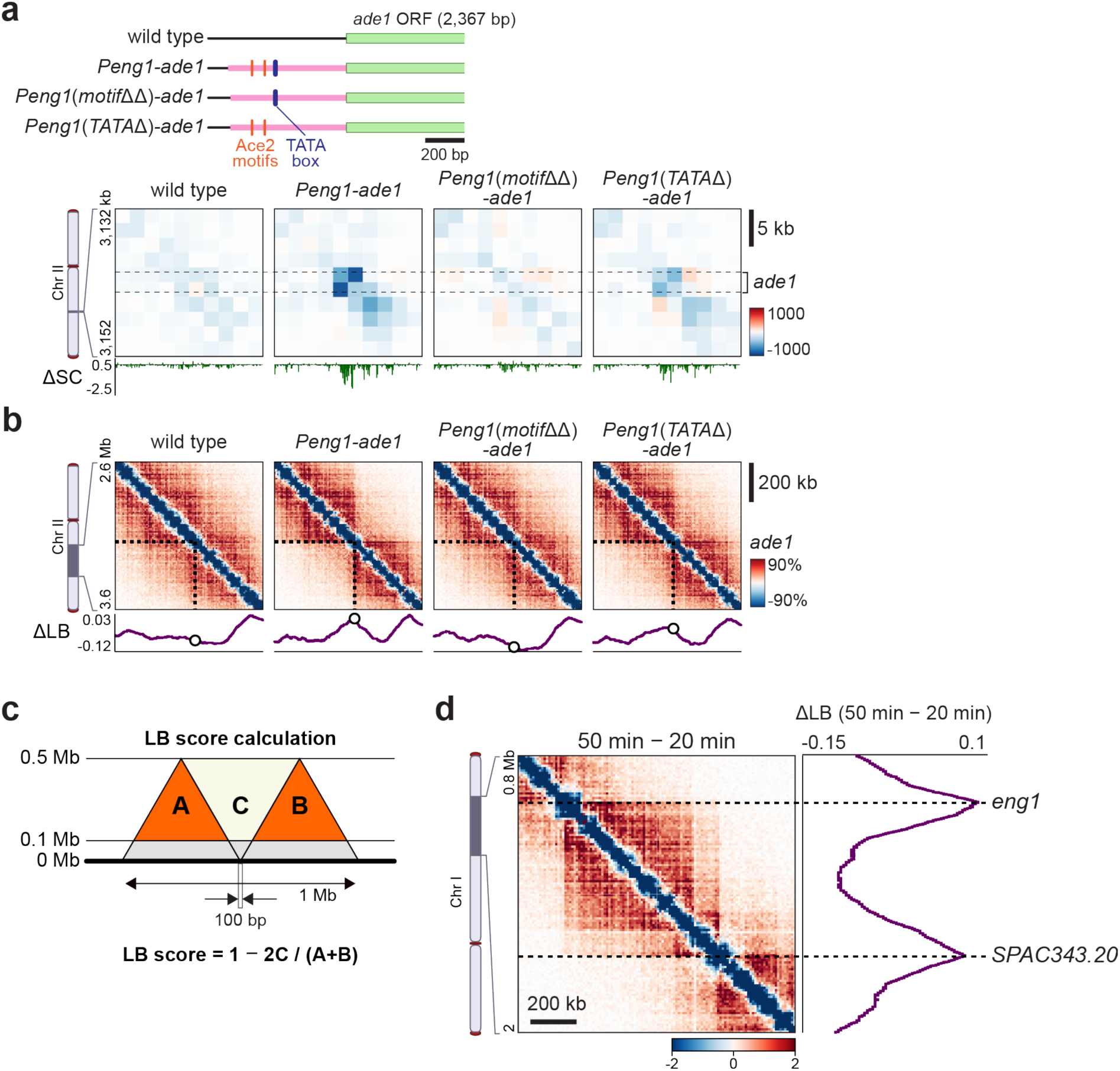
Local decompaction is tightly associated with boundary formation. **a**, Impaired decompaction in the *Peng1* mutants. The top panel illustrates the *ade1* gene locus in the wild-type and the *Peng1-ade1* strain series. Differential contacts between 20 and 50 minutes in each strain are shown in the bottom panels. Below each Hi-C difference map is its ΔSC score plot in the same region. **b**, Defective boundary formation with the mutant *Peng1*. The difference maps, as in Fig. 3a, are shown for the 1-Mb region of chromosome II that contains the *ade1* gene. Below each Hi-C difference map is its ΔLB score plot (see also Fig. 3c**,d**). **c**, Calculation schemes of long-range boundary (LB) scores and differential LB (ΔLB) scores between 20 and 50 minutes after G2/M block release. **d**, A ΔLB score plot at the example locus that contains the *eng1* and *SPAC343.20* genes. The same difference map as Fig. 1b but at a lower resolution (10-kb bins) is shown, and adjacent to it is its ΔLB score plot.

To verify the above observations systematically and quantitatively, we sought to develop a computational method to quantify a boundary activity at a given locus. To this end, we defined a long-range boundary (LB) score for each 100-bp bin, as in **Fig. 3c**. As a validation, we calculated LB scores based on the Hi-C results at 20 and 50 minutes after G2/M block release shown in **Fig. 1b** and found that the relative LB scores between these two time points (ΔLB scores) peaked at the *eng1* and *SPAC343.20* gene loci (**Fig. 3d**), which is consistent with the previous study that identified prominent domain boundaries at these gene loci during mitosis^19^. We calculated the LB scores around the *ade1* gene based on the Hi-C results of the *Peng1-ade1* strain series shown in **Fig. 3b**. Compared to the wild-type background without the *Peng1* insertion, the *Peng1* insertion resulted in higher LB scores at the *ade1* locus. (**Fig. 3b**). By contrast, this ΔLB score at the *ade1* locus was comparable between wild-type and *Peng1*(*motif*ΔΔ)*-ade1* cells, and the *Peng1*(*TATA*Δ) insertion showed a weaker boundary activity compared to the wild-type *Peng1* insertion (**Fig. 3b**). These quantitative comparisons further support the tight relationship between local decompaction and boundary formation.

### Local decompaction-inducible promoters can form a long-range contact boundary

We expanded the quantification of boundary formation, LB scoring, to the entire fission yeast genome using the Hi-C results of mitotically synchronized cells to examine whether the hyper-decompaction genes are associated with boundary formation. As a result, we found a coincidence of intense local decompaction (negative ΔSC score peaks) and robust boundary formation (positive ΔLB score peaks) at some gene loci (**Extended Data Fig. 5a**). Statistically, however, the two scores did not show a good correlation (**Extended Data Fig. 5b**), even though we have established so far that these two events are closely connected. One obvious explanation is that, while the ΔSC scores show sharp peaks at the hyper-decompaction genes, the ΔLB peaks are much broader, spanning hundreds of kilobases, which can render the ΔLB scores of many genes high when they are located near the ΔLB peak positions even if they do not have the boundary forming activity. Another possibility is that the decompaction-linked boundary formation is only active in some specific chromatin contexts, as in the previous observations that random chromosome insertions of the TAD boundary-competent DNA fragment result in variable outcomes depending on the sites of insertion^26,27^. To overcome these potential problems in assessing boundary-forming activities, we made use of the fact that *Peng1* can trigger boundary formation when ectopically placed at *ade1* (**Fig. 2g**), which means that the chromatin state around the *ade1* gene locus is permissive to boundary formation. Therefore, we picked several genes with various ΔSC scores, Ace2 binding, and mitotic expression status (**Fig. 4a**), and a promoter region from each gene (**Extended Data Fig. 6a**) was inserted at the *ade1* gene as in the *Peng1-ade1* strain to test its ability to establish the *de novo* boundary. Here, we examined Ace2-bound promoters of the *cfh4* and *adg3* genes (*Pcfh4* and *Padg3*; gene ΔSC scores, -0.531 and -0.100, respectively, see also **Extended Data Fig. 6b)** as well as *Peng1* (gene ΔSC score, -0.744), and other mitotically activated promoters of the *cdc22*, *cdc18*, and *tos4* genes (*Pcdc22*, *Pcdc18*, and *Ptos4*; gene ΔSC scores, -0.354, -0.095, and -0.167, respectively), at which Ace2 enrichment was negligible or not detected (**Fig. 4a**). As a comparison, a constitutively and highly active promoter of the *adh1* gene (*Padh1*; gene ΔSC score, -0.147) was also tested (**Fig. 4a**). After verifying that the *ade1* gene was indeed subject to mitotic activation in the presence of these ectopic promoters except for *Padh1* (**Extended Data Fig. 6c**), we performed Hi-C in mitotically synchronized cells in these modified *ade1* gene backgrounds and calculated the ΔSC and ΔLB scores of the *ade1* gene for each strain (**Fig. 4b,c**). Among the Ace2-bound promoters tested here, *Pcfh4*, which has a strong decompaction activity at its endogenous locus (**Fig. 4a**), was moderately capable of decompaction and boundary formation at *ade1* (**Fig. 4b,c**) with the gene ΔSC and gene ΔLB scores at *ade1* (denoted hereafter as ΔSC[*ade1*] and ΔLB[*ade1*], respectively) being -0.381 and -0.012, slightly less efficient than with *Peng1* (ΔSC[*ade1*]: -0.681, ΔLB[*ade1*]: 0.015). Surprisingly, however, the other promoter targeted by Ace2 tested here, *Padg3*, which does not display intense decompaction at its endogenous site (**Fig. 4a**), was not efficient in decompaction nor boundary formation at the ectopic site as well (**Fig. 4b,c**; ΔSC[*ade1*]: -0.258, ΔLB[*ade1*]: -0.059). Furthermore, even though no apparent Ace2 binding was detected at the endogenous *cdc22* gene in our Ace2 ChIP-seq data (**Fig. 4a**), *Pcdc22* showed a strong activity in local decompaction and large-scale boundary formation when placed at *ade1* (**Fig. 4b,c**; ΔSC[*ade1*]: -0.520, ΔLB[*ade1*]: 0.039), indicating that other mechanism(s) than the Ace2-driven mitotic activation should operate at *Pcdc22*. Indeed, the *cdc22* gene is known to be activated by the MluI binding factor (MBF) transcription factor complex^28^, and we also verified the binding of the Cdc10 subunit of MBF to the endogenous *cdc22* promoter by ChIP-seq (**Extended Data Fig. 7a-c**). To explore the possible involvement of MBF, we constructed a mutant derivative of *Pcdc22*, referred to as *Pcdc22*(*MluIΔΔΔ*), that lacks the three MluI restriction sites (5′-ACGCGT-3′ x3), which are putative MBF binding sites^28^. After verifying that these elements were indeed necessary for mitotic gene activation (**Extended Data Fig. 7d**), we performed Hi-C and found weakened boundary formation and decompaction activities of the mutant *Pcdc22* compared to the wild-type counterpart (**Extended Data Fig. 7e,f**; ΔSC[*ade1*]: - 0.144, ΔLB[*ade1*]: -0.052 in case of *Pcdc22*(*MluIΔΔΔ*); ΔSC[*ade1*]: -0.520, ΔLB[*ade1*]: -0.039 in case of wild-type *Pcdc22*), suggesting that the MBF transcription factor can exert a regulatory role in short- and long-range DNA contacts as Ace2 can do. Interestingly, *Pcdc18* and *Ptos4*, which are also targeted by the MBF transcription factor (**Extended Data Fig. 7b,c**), were not effective in boundary formation (**Fig. 4c**; ΔLB[*ade1*]: -0.065 in case of *Pcdc18*; ΔLB[*ade1*]: -0.058 in case of *Ptos4*), and these promoters poorly induced local decompaction at the ectopic site (**Fig. 4b**; ΔSC[*ade1*]: -0.152 in case of *Pcdc18*; ΔSC[*ade1*]: -0.204 in case of *Ptos4*) as well as at its endogenous site (**Fig. 4a**; ΔSC[*cdc18*]: -0.095, ΔSC[*tos4*]: -0.167), indicating that MBF binding is not sufficient for boundary formation. Gathering all these Hi-C data of the ectopic promoters at *ade1* including the mutant derivative data in **Fig. 3a,b** and **Extended Data Fig. 7e,f**, we tested a possible correlation between local decompaction and boundary formation activities at this locus. Intriguingly, the ΔSC and ΔLB scores at the *ade1* gene loci with various ectopic promoters showed a strong negative correlation between these two scores (**Fig. 4d**; Pearson correlation coefficient: *r* = -0.881). Taken together, local decompaction is, among others, the most critical determinant of boundary formation, raising the possibility that intense local decompaction can drive the formation of long-range contact boundaries.

**Fig. 4.**
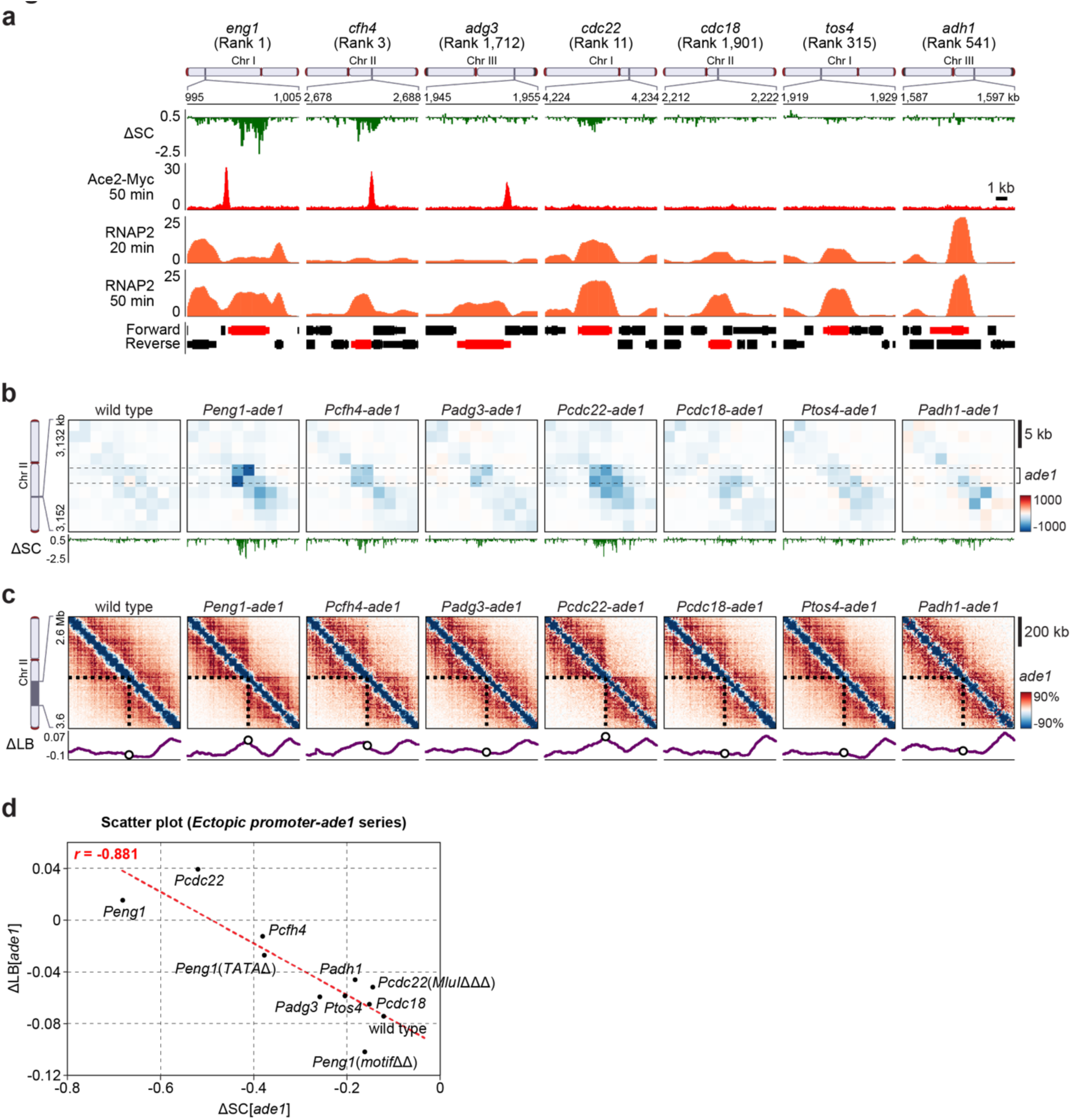
Ectopic promoter experiments reveal local decompaction activity best correlates with boundary formation competency. **a**, Gene promoters used for the ectopic insertion experiments in this study. The genes shown in this figure were selected so that a variety of patterns in Ace2 binding (Ace2-Myc ChIP-seq), ΔSC scores, and mitotic gene activation (RNAP2 ChIP-seq) could be examined. **b**, Decompaction activities of the ectopic promoters inserted into the *ade1* gene. Differential contacts between 20 and 50 minutes in each strain are shown for the 20-kb region containing the *ade1* gene, and below each Hi-C map is its ΔSC score plot. **c**, Boundary formation activities of the ectopic promoters inserted into the *ade1* gene. Differential contacts between 20 and 50 minutes in each strain are shown for the indicated 1-Mb region of chromosome II, and below each Hi-C map is its ΔLB score plot. **d**, Examination of a possible correlation between local decompaction and boundary levels. A scatter plot of the gene ΔSC and gene ΔLB scores of *ade1* (ΔSC[*ade1*] and ΔLB[*ade1*], respectively) in the wild-type and ectopic promoter insertion strains is shown. The trend line is overlaid as a red dotted line. A Pearson correlation coefficient, *r*, was calculated to assess the possible correlation between the two scores.

### Local decompaction and boundary formation efficiencies are dependent on length of transcriptional elongation

The fact that the local hyper-decompaction predominantly occurs within transcribed regions (**Extended Data Fig. 3a**) led to the question as to whether transcriptional elongation plays a role in this local chromatin regulation. To address this question, we set out to construct strains with an ectopic transcription terminator sequence insertion at *eng1* so that transcription can be terminated prematurely. For this purpose, we utilized the terminator element of the *CYC1* gene in budding yeast, which was previously reported to be functional even in fission yeast^29^. We picked four positions within the transcribed region of the *eng1* gene and inserted the *CYC1* terminator (*T_CYC1_*) so that the distance of *eng1* transcription was to be 1 bp, 1,086 bp, 2,247 bp, or 3,369 bp (**Fig. 5a**).

**Fig. 5.**
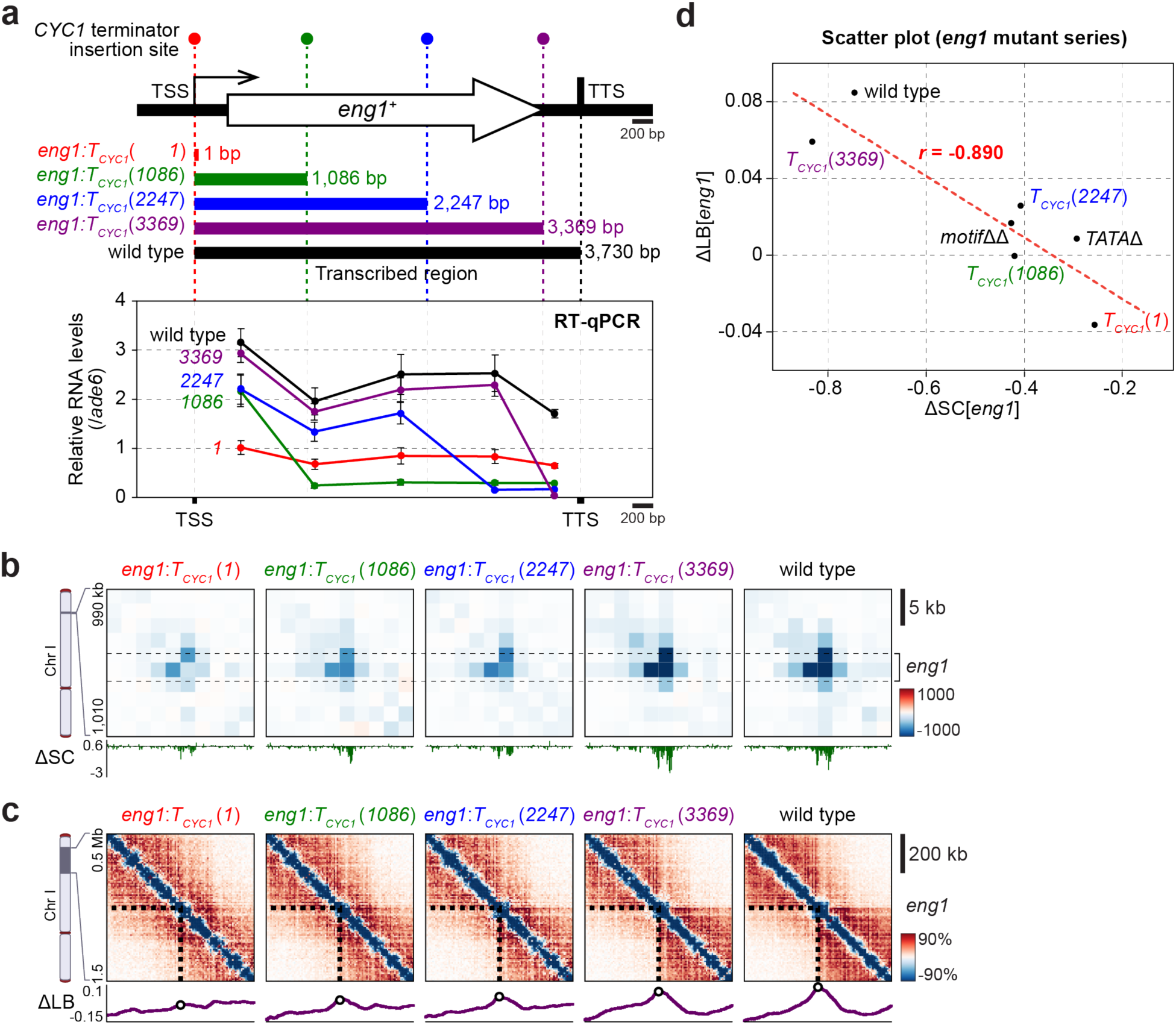
Transcriptional elongation is necessary for efficient local decompaction and boundary formation. **a**, A schematic of terminator insertion strains on the top. Four positions indicated by dots and dotted lines in red, green, blue, and purple that are located at 1 bp, 1,086 bp, 2,247 bp, and 3,369 bp downstream of the transcription start site (TSS) were selected as insertion sites of the transcriptional terminator sequence of the budding yeast *CYC1* gene (*T_CYC1_*) to generate four termination insertion strains (*eng1*:*T_CYC1_*). Note that the distance between TSS and the transcription termination site (TTS) of the *eng1* gene in wild type is 3,730 bp. Below is the result of RT-qPCR to verify the expected transcription termination by the *T_CYC1_* insertion. Mitotically synchronized *cdc25-22* cells in the wild-type and terminator insertion backgrounds were harvested at 50 minutes after release from the G2/M transition, and RT-qPCR was performed. The RT-qPCR primer positions exactly correspond to the upper schematic of the *eng1* gene. The RNA levels relative to that of *ade6* were shown, and the error bars represent the standard error of the mean (n=3). **b**, Defective local decompaction at the *eng1* locus by early transcriptional termination. Differential contacts between 20 and 50 minutes in each strain are shown for the 20-kb region containing the *eng1* gene, and below each Hi-C difference map is its ΔSC score plot. **c**, Defective boundary formation at the *eng1* locus by early transcriptional termination. Differential contacts between 20 and 50 minutes in each strain are shown for the indicated 1-Mb region of chromosome I, and below each Hi-C difference map is its ΔLB score plot. **d**, Examination of a possible correlation between local decompaction and boundary levels. A scatter plot of the gene ΔSC and ΔLB scores of *eng1* (ΔSC[*eng1*] and ΔLB[*eng1*]) in the wild-type and terminator insertion strains is shown. The trend line is overlaid as a red dotted line. A Pearson correlation coefficient, *r*, was calculated to assess the possible correlation between the two scores.

After verifying the expected termination of *eng1* transcription in the *T_CYC1_* insertion strains (*eng1*:*T_CYC1_*) by RT-qPCR (**Fig. 5a**), we performed Hi-C and found that the robust local decompaction at *eng1* requires sufficiently long transcription distance (**Fig. 5b**). Interestingly, the boundary formation activity became stronger with the longer elongation (**Fig. 5c**). Gathering all the Hi-C data of the *eng1* mutants examined in **Fig. 2c,f** and the terminator insertion strains *eng1*:*T_CYC1_* examined in **Fig. 5b,c**, we again found a strong correlation between ΔSC and ΔLB scores at the *eng1* gene (**Fig. 5d**; Pearson correlation coefficient: *r* = -0.890), further supporting the notion that local decompaction is critical for boundary formation.

### Local decompaction-competent promoters are active in histone removal

Our terminator insertion experiments in **Fig. 5** suggest that transcriptional elongation during mitosis can render the local chromatin highly decompacted. Given that active transcription is associated with nucleosome depletion at promoters and gene bodies^30^, we next asked whether the hyper-decompaction genes undergo histone loss during mitosis. We performed ChIP-seq for canonical histone H3 using mitotically synchronized cells at 20 and 50 minutes after G2/M block release and found that the histone H3 occupancy was strongly diminished upon mitosis at only tens of gene loci across the genome, such as the *eng1* and *SPAC343.20* genes (**Fig. 6a**). Surprisingly, mitotic gene activation is not always associated with intense histone eviction. Of the genes whose promoter sequence was used for the ectopic promoter strains in **Fig. 4**, the *eng1* and *cdc22* genes underwent strong histone H3 loss during mitosis, but the *adg3* and *cdc18* genes did to much lesser extent, even though these genes are significantly up-regulated during mitosis (**Fig. 6b**). Given that the former two genes are subject to local decompaction during mitosis but the latter two are not (**Fig. 6b**), these histone H3 ChIP results raise the possibility that histone depletion is tightly connected to local decompaction. Of note, however, the decompaction-competent *cfh4* gene was subject to the same level of histone H3 loss as the decompaction-incompetent *tos4* gene. Consistently, the levels of histone H3 loss and local decompaction during mitosis at all the protein-coding genes did not show good correlation (**Fig. 6c**). Nevertheless, we still observed that the hyper-decompaction genes are highly enriched in genes with strong histone loss (**Fig. 6c**), suggesting that histone removal during mitosis can just promote, but is not the same phenomenon as, intense local decompaction. To further support this observation, we performed a histone H3 ChIP analysis of the *eng1* mutants, *eng1-TATA*Δ and *eng1-motif*ΔΔ, and found that the mitotic histone removal was suppressed by the mutations (**Fig. 6d**); in the *eng1-TATA*Δ mutant, histone H3 remained associated with the *eng1* ORF, even though histone H3 was lost at the *eng1* promoter to the same extent as in the wild-type strain; in the *eng1-motif*ΔΔ, the histone H3 eviction was suppressed both at the promoter and ORF. Since these mutations cause defective local decompaction (**Fig. 2c**), this histone occupancy analysis suggests that, at least at the *eng1* gene locus, intense decompaction is promoted by histone depletion that is dependent on Ace2-mediated gene up-regulation during mitosis.

**Fig. 6.**
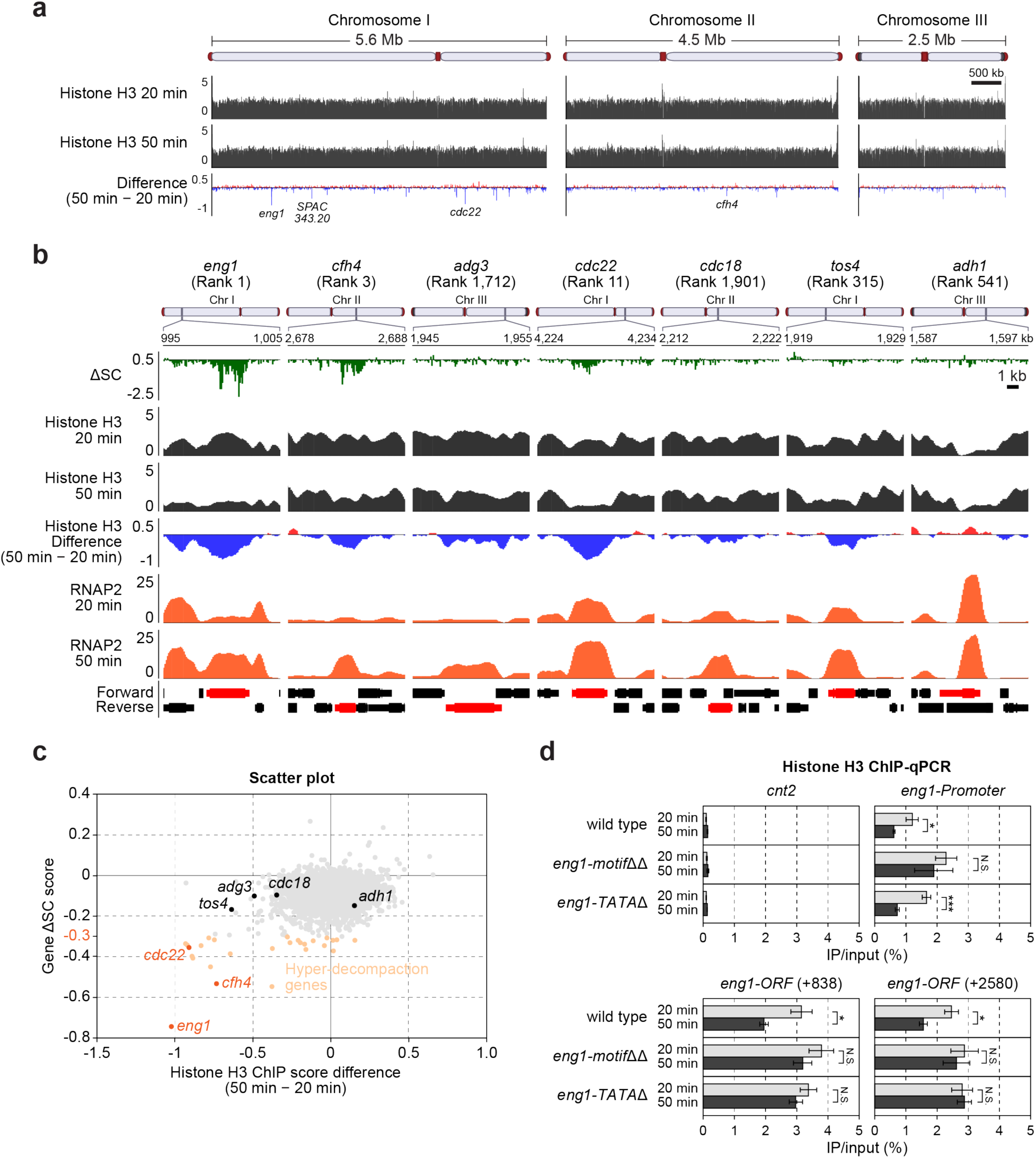
Mitotic histone eviction can support local decompaction. **a,** Genome-wide binding profiles of histone H3 before and during mitosis. Mitotically synchronized *cdc25-22* cells were fixed at 20 and 50 minutes after release from the G2/M block, and ChIP-seq was performed using anti-histone H3 antibody. The difference between 20 and 50 minutes was calculated, and the increased and decreased ChIP-seq signals over time are indicated by red and blue colors, respectively. **b,** Histone eviction activities at select genes, the promoters of which were used for the ectopic promoter strains. The RNAP2 ChIP-seq plots at 20 and 50 minutes and the ΔSC score plots are shown in addition to the histone H3 ChIP-seq data to explore the possible correlation between gene activation, local decompaction activity, and histone eviction. **c**, Examination of a possible correlation between histone depletion and decompaction levels. A scatter plot of the differential histone H3 ChIP-seq scores between 20 and 50 minutes and the gene ΔSC scores is shown. **d**, Defective histone eviction at the *eng1* gene in the *eng1-TATA*Δ and *eng1-motif*ΔΔ mutants. Mitotically synchronized *cdc25-22* cells were fixed at 20 and 50 minutes after release from the G2/M block, and ChIP-qPCR was performed using anti-histone H3 antibody. The positions of the qPCR primer sets designed in the open reading frame (ORF) of the *eng1* gene are indicated by the base-pair distance between the middle of the PCR amplicon and the initial codon. The core centromere region of chromosome II, *cnt2*, at which canonical histone H3 is replaced by the CENP-A variant, was analyzed as a negative control locus. DNA recovery efficiencies after immunoprecipitation relative to input DNA (IP/input) are shown, and the error bars represent the standard error of the mean (n=3). Statistical significance was assessed using a two-tailed *t*-test (N.S., *P* > 0.05; *, *P* < 0.05; ***, *P* < 0.001).

## DISCUSSION

In this study, by mapping fine-scale chromosomal DNA contacts in mitotic fission yeast cells with a modified Hi-C approach (**Extended Data Fig. 1**), we discovered that the intense local chromatin decompaction at a limited number of gene loci in the context of large-scale chromosomal reorganization during mitosis (**Fig. 1**). This finding is consistent with previous microscopic observation: Cryo-electron tomography of fission yeast mitotic chromosomes detected unevenly condensed chromatin with a small portion of open “pockets,” which are free of macromolecular complexes^13^. Moreover, our data suggest that this small-scale decompaction of chromatin can be facilitated by local histone eviction, which requires the binding of specific transcription factors to promoters of the mitotically activated genes followed by transcriptional activation (**Fig. 6d** and **Extended Data Fig. 4a**). One might assume, however, that this mitotic local decompaction is simply attributed to up-regulated transcription, because previous high-resolution mappings of nucleosomal DNA contacts using micrococcal nuclease have demonstrated that levels of gene-sized, local DNA compaction are anti-correlated with transcriptional activity in yeast^31,32^. Contrary to this notion, our ectopic promoter analyses in **Fig. 4** show that Ace2-driven gene activation is not always sufficient for effective local decompaction, indicating that this local event is not a simple outcome of increased transcription levels. Importantly, this small-scale decompaction of local chromatin seems to be tightly connected to the formation of boundaries that demarcate large-scale mitotic chromosomal domains. Even though we demonstrated that binding of the Ace2 transcription factor to the *eng1* promoter is necessary for the efficient formation of the boundary at this gene locus (**Fig. 2**), our ectopic promoter analyses in **Fig. 4** showed that Ace2-driven gene activation is not always sufficient for chromosomal domain demarcation during mitosis. Interestingly, the results in **Fig. 4** also led to the discovery that another mitotically engaged transcription factor, MBF, has a potential in boundary formation, but again, mere gene activation by MBF cannot account for demarcation of mitotic chromosome domains. Instead, our data demonstrated that chromatin compaction level shows a robust negative correlation with boundary strength (**Fig. 4d and 5d**), leading to the novel concept that local chromatin decompaction can be a critical contributor to the organization of the mitotic chromosome structure (**Extended Data Fig. 8a**).

Furthermore, our results point to the possible molecular mechanism of the local decompaction-mediated boundary formation. By analyzing the terminator insertion strain series, we found that the length of transcriptional elongation at the *eng1* gene is critical for efficient boundary formation at this locus (**Fig. 5**), which raises the question of how elongation distance contributes to the insulation of domain contacts. A simple explanation is that a stretched chromatin fiber induced by local decompaction upon transcription elongation can provide a sufficient physical distance between the two adjoining domains, thereby preventing them from associating with each other. However, this idea disregards the chromosome elasticity that allows for contacts between the two domains by, for example, looping out the flexible chromatin in between. Another possibility is based on the transcription-driven DNA motion: DNA can rotate along its helical axis when transcription machinery runs through it^33^. It might be thus attractive to speculate that more extended transcription elongation can drive more rotary motion of two domains located upstream and downstream so that their stable contacts are hampered. In this case, decompaction may provide the local chromatin with enough flexibility, enabling efficient transmission of the rotary motion to surrounding chromatin domains (**Extended Data Fig. 8b**).

Another potential impact of this study is that our findings here can be extended to a better understanding of the principle of chromosomal DNA organization, even in higher eukaryotes. In general, topologically associated domains (TADs) are assembled predominantly by another SMC protein-containing complex, cohesin^34^, and similar to condensin-mediated domains in fission yeast mitosis, TAD boundaries can be established in a transcription-dependent manner^26,27,35,36^. However, it remains obscure why active transcription does not always associate with TAD boundary formation. Our study indicated that specific transcription factors have a potential in boundary formation and, more importantly, that intense local decompaction can constitute an additional regulatory layer. In the future, it would be interesting to test whether the formation of TAD boundaries in higher eukaryotes also requires specific transcription factors and local decompaction.

We have previously demonstrated that boundaries of condensin-mediated long-range contacts are critical for faithful segregation of mitotic chromosomes^7^. In the present study, we have provided ample experimental data on how a subset of mitotically activated genes, by exerting their ability in local chromatin decompaction, can help organize the mitotic chromosome configuration. In this context, the *SPAC343.20* gene, one of the hyper-decompaction genes (**Fig. 1c,h**), encodes a small peptide with only 113 amino acids, but its transcription extends to 2 kb and overlaps with a convergently arranged gene, shedding light on how fission yeast utilizes this unnaturally extended gene as a tool to organize the functional chromatin domain required for domain-based chromosomal compaction and segregation. Therefore, our research thus raises an intriguing possibility that the distribution of mitotically activated genes along chromosomes and their length have been optimized during evolution. From this point of view, we expect that our findings also provide an important hint about the synthetic biology approach, where artificially synthesized genomes are well-designed so that they can be stably maintained over generations.

## EXPERIMENTAL PROCEDURES

### Fission yeast strains and culture conditions

All experiments in this study were performed using fission yeast *Schizosaccharomyces pombe* strains listed in **Supplementary Table 2**. Strain constructions were carried out using the standard chemical transformation and genetic cross approaches. A DNA construct for epitope tagging to the *cdc10* gene was amplified by polymerase chain reaction (PCR) as described previously^37^. The ectopic promoter strains (**Fig. 4**) were also constructed using a similar PCR-based approach. The Ace2 binding sites and TATA box were excised from the *eng1* promoter without leaving any selectable markers by two rounds of transformation: first, the *ura4^+^* gene integration at the *eng1* promoter; second, *ura4^+^* removal with leaving the deletion. This markerless mutagenesis strategy was also applied to inserting the *CYC1* terminator sequence (*T_CYC1_*) in **Fig. 5**. The PCR primers used for the strain constructions in this study are listed in **Supplementary Table 3**.

Cell cycle synchronization was performed using the *cdc25-22* temperature-sensitive mutant allele. Cells harboring this mutation were grown to logarithmic phase (OD_595_ ∼0.25) in yeast extract-adenine (YEA) liquid medium at the permissive temperature (26°C) and further cultured at the restrictive temperature (36°C) for 3.5 hours so that the cells were arrested at the G2/M transition. The cultures were quickly cooled in an ice-water bath, allowing synchronous release into the M phase at 26°C. For experiments with the *cut14-208* condensin and *rad21-K1* cohesin temperature-sensitive mutant strains^15,38^, cells were grown to logarithmic phase (OD_595_∼0.4) in YEA liquid medium at the permissive temperature (26°C) before incubation at the restrictive temperature (36°C) for 1 hour. For the other experiments with asynchronous cultures, cells were grown to logarithmic phase (OD_595_ ∼0.5) in YEA liquid medium at 30°C.

### High-resolution in situ Hi-C

In situ Hi-C experiments were performed as described previously^19^ with some modifications in cell fixation and DNA digestion conditions to obtain Hi-C maps at a higher resolution than before (see also **Extended Data Fig. 1** and **Supplementary Note**). A total of 4×10^8^ cells were fixed with 3% paraformaldehyde (pFA; Sigma, P6148) at the culture temperature for 10 minutes and disrupted with Mini-Beadbeater-16 (BioSpec Products) in the presence of lysis buffer [50 mM HEPES (Fisher BioReagents, BP310) pH 7.5, 140 mM NaCl (Sigma, S6191), 1 mM ethylenediaminetetraacetate dihydrate (EDTA; Invitrogen, 15575-038), 1% Triton X-100 (Sigma, T8787), 0.1% sodium deoxycholate (Sigma, D6750), 1x Complete (Roche, 11836170001), and 1 mM phenylmethanesulfonyl fluoride (PMSF; Sigma, P7626)] and glass beads (Sigma, G8772). The disrupted cells were further fixed with 3 mM disuccinimidyl glutarate (DSG; Thermo Scientific, 20593) at 30°C for 40 minutes. The dually fixed cells were successively treated with 0.1% sodium dodecyl sulfate (SDS; Invitrogen, 15553-035) at 62°C for 7 minutes, with 1% Triton X-100 at 37°C for 15 minutes, and with the MboI (New England Biolabs, R0147), HinfI (New England Biolabs, R0155), and MluCI (New England Biolabs, R0538) restriction enzymes (0.1 U/µL for each enzyme) at 37°C overnight to fragment genomic DNA. After inactivation of the enzymes at 62°C for 20 minutes, the protruding ends generated by the restriction digestion were filled in by 0.13 U/uL Klenow fragment (New England Biolabs, M0210) treatment at 37°C for 45 minutes in the presence of 50 µM biotinylated deoxyadenosine triphosphate (biotin-14-dATP; Jena Bioscience, NU-835-BIO14) in addition to other deoxyribonucleotides [dTTP (Invitrogen, 55085), dGTP (Invitrogen, 55084), and dCTP (Invitrogen, 55083); 500 µM for each] to generate blunt ends with biotin label. The blunt ends were subjected to proximity ligation by 4.17 U/µL T4 DNA ligase (New England Biolabs, M0202) treatment at room temperature for 4 hours. After crosslinks were reversed by incubating at 65°C for 4 hours in the presence of 200 mM NaCl, 0.5% SDS, and 500 µg/mL proteinase K (Invitrogen, 25530049), DNA was purified by Tris-EDTA-saturated neutral (pH 8.0) phenol:chloroform:isoamyl alcohol (neutral PCI; Sigma, P2069) extraction and ethanol precipitation. The DNA was fragmented by sonication using Bioruptor UCD-200 (Diagenode) and further purified with 1× volume of AMPure XP (Beckman Coulter, A63881). The biotinylated DNA was selectively pulled down using Dynabeads MyOne Streptavidin T1 (Invitrogen, 65601) magnetic beads at 25°C for 15 minutes and subsequently subjected to DNA-end repair with 0.5 U/µL T4 polynucleotide kinase (New England Biolabs, M0201), 0.12 U/µL T4 DNA polymerase (New England Biolabs, M0203), and 0.05 U/µL Klenow fragment at 25°C for 30 minutes, followed by dA tailing with 0.25 U/µL Klenow (3′→5′ exo-) (New England Biolabs, M0212) in the presence of 0.5 mM dATP (Invitrogen, 55082) at 37°C for 30 minutes. The DNA ends with dA overhangs were ligated to NEBNext Adaptor for Illumina (New England Biolabs) using NEBNext Ultra II Ligation Module (New England Biolabs, E7595) and USER Enzyme (New England Biolabs), according to the manufacturer instructions. The adaptor-ligated DNA on the magnetic beads was amplified by 5-8 cycles of PCR using NEBNext Ultra II Q5 Master Mix (New England Biolabs, M0544) and NEBNext Multiplex Oligos for Illumina (New England Biolabs). The PCR-amplified libraries were purified with 0.9× volume of AMPure XP before sequencing runs with NextSeq 500, NextSeq 2000, or NovaSeq 6000 (Illumina) to obtain paired-end reads.

### In situ Hi-C data processing

In situ Hi-C data was processed as described previously using the rfy_hic2 packages (https://github.com/rafysta/rfy_hic2)^19^. Briefly, paired-end reads from next-generation sequencing were aligned separately to the *S. pombe* genome (2018) using an iterative alignment strategy with bowtie2 (version 2.4.4). Redundant paired reads derived from PCR bias, reads aligned to repetitive sequences, and reads with low mapping quality (MapQ < 30) were removed. Read pairs with inward or outward orientations within 10 kb were excluded from the analysis, as they contain undigested products or self-ligation artifacts. The number of correct Hi-C reads in inward and outward orientations was estimated using the data of reads aligned in the same orientation. The fission yeast genome was divided into non-overlapping bins of 2 kb and 10 kb to create raw contact matrices by counting the number of read pairs assigned to respective bin combinations. When assigning reads to bins, the original fragments, which were cut by one to three restriction enzymes, were determined from the aligned positions, and the reads were assigned to the bin containing the center of the original fragment. All contact maps used in this study were generated from raw contact maps. Read numbers before and after the respective filtering processes were summarized in **Supplementary Table 4**.

### Short-range compaction (SC) scores

The schematic representations of SC score calculation are shown in **Fig. 1d** and **f**. Paired-end sequence reads with MapQ > 30 that were aligned in the same orientation (i.e., correct Hi-C reads that do not include self-ligation or undigested products) were extracted if the alignment distance between the two reads was less than 2 kb. The entire chromosome was divided into 100 bp segments, and a score was calculated for each segment. After the total number of reads for each 100 bp segment was calculated, this score was divided by the total number of reads, and the result was multiplied by 125718.2, which is the total length of the *S. pombe* genome (12,571,820 bp) divided by 100. This calculation yielded the SC score, with an expected value of 0.01. The ΔSC score was calculated simply by subtracting the 20-minute SC score from the 50-minute SC score for each bin, and the gene ΔSC score was determined by averaging the ΔSC scores of bins that overlap the transcribed region of the gene of interest as in **Fig. 1f**.

### Long-range boundary (LB) scores

The schematic representation of LB score calculation is shown in **Fig. 3c**. Paired-end sequence reads with MapQ > 30 that were aligned in the same orientation (i.e., correct Hi-C reads that do not include self-ligation or undigested products) were extracted if the alignment distance between the two reads was between 100 kb and 500 kb. The entire chromosome was divided into 100 bp segments, and a score was calculated for each segment. The score was calculated by defining the genomic contacts between the upstream and downstream regions of the target bin as “inter” and the contacts within the upstream and downstream regions as “intra.” After calculating the total number of reads, the LB score at the target bin was defined as 1 - (inter × 2 / intra). The ΔLB score was calculated simply by subtracting the 20-minute LB score from the 50-minute LB score for each bin, and the gene ΔLB score was determined by averaging the ΔLB scores of bins that overlap the transcribed region of the gene of interest in a similar manner to the gene ΔSC score calculation (**Fig. 1f**).

### Chromatin immunoprecipitation (ChIP)

ChIP experiments were performed as described previously^39^ with minor modifications. A total of 4×10^8^ cells were first fixed with 3% pFA at the culture temperature for 10 minutes and subsequently with 10 mM dimethyl adipimidate (DMA; Sigma, 285625) at room temperature for 45 minutes. The fixed cells were disrupted with Mini-Beadbeater-16 in the presence of lysis buffer (50 mM HEPES pH 7.5, 140 mM NaCl, 1 mM EDTA, 1% Triton X-100, 0.1% sodium deoxycholate, 1x Complete, and 1 mM PMSF) and glass beads. The crude extract was subject to sonication with Bioruptor UCD-200 so that DNA was fragmented to approximately 300 bp on average. After clarification of the crude extract by centrifugation, the lysate was incubated at 4°C for 2 hours with Dynabeads Protein G (Invitrogen, 10003D) magnetic beads that bound anti-RNA polymerase II C-terminal domain antibody CTD4H8 (Abcam, ab5408), anti-Myc antibody 9B11 (Cell Signaling Technology, 2276), or anti-Pk antibody SV5-Pk1 (Bio-Rad, MCA1360). The magnetic beads after the immunoprecipitation were washed once with Buffer 1 (50 mM HEPES pH 7.5, 140 mM NaCl, 1 mM EDTA, 1% Triton X-100, and 0.1% sodium deoxycholate), once with Buffer 1ʹ (50 mM HEPES pH 7.5, 500 mM NaCl, 1 mM EDTA, 1% Triton X-100, and 0.1% sodium deoxycholate), once with Buffer 2 [10 mM Tris pH 8.0 (Invitrogen, 15568-025), 250 mM lithium chloride (Sigma, L4408), 1 mM EDTA, 0.5% IGEPAL (Sigma, I8896), and 0.5% sodium deoxycholate], and twice with TE buffer (10 mM Tris pH 8.0 and 1 mM EDTA). The washed beads were treated with 10 µg/mL RNase A (Thermo Scientific, EN0531) at 37°C for 15 minutes, and the immunoprecipitates were eluted twice by incubating the beads in TE buffer containing 1% SDS at 65°C for 15 minutes. After crosslinks were reversed by incubating at 65°C overnight in the presence of 200 mM NaCl and 250 µg/mL proteinase K (Invitrogen, 25530049), co-precipitated DNA was purified with QIAquick PCR Purification Kit (Qiagen, 28104) or by neutral PCI extraction and ethanol precipitation. In a similar fashion, the lysate was treated with RNase A and Proteinase K, and DNA purification was performed to obtain input DNA. The immunoprecipitated DNA and input DNA were analyzed by quantitative PCR (qPCR) with Fast SYBR Green Master Mix (Applied Biosystems, 4385612) and StepOnePlus Real-Time PCR System (Applied Biosystems). The qPCR primers used in this study are listed in **Supplementary Table 3**. To prepare a ChIP sequencing (ChIP-seq) library, the immunoprecipitated DNA was subjected to end repair and adaptor ligation reactions using NEBNext Ultra II DNA Library Prep Kit for Illumina (New England Biolabs, E7645), according to the manufacturer instruction. The adaptor-ligated DNA was amplified by 5-12 cycles of PCR using NEBNext Ultra II Q5 Master Mix and NEBNext Multiplex Oligos for Illumina. The PCR-amplified libraries were purified with 0.9× volume of AMPure XP before sequencing runs with NextSeq 500, NextSeq 2000, and NovaSeq 6000 to obtain paired-end reads.

### ChIP-seq data processing

Paired-end sequenced reads were aligned to the *S. pombe* genome (2018) using Bowtie2 (version 2.4.4) in paired-end mode. Read pairs that were correctly oriented and aligned at the appropriate distance as paired-end reads were extracted using Samtools (version 1.19) with the option -f 0x2. BigWig files were generated using the bamCoverage function of deepTools (version 3.5.4). To minimize the influence of rDNA repeats, normalization was performed to achieve an average score of 1, excluding chromosome III, which contains rDNA repeats. For this purpose, the specific options used were “--ignoreForNormalization III --normalizeUsing RPGC --effectiveGenomeSize 10118937.” For the RNAP2, Cut14-Pk, and histone H3 ChIP-seq, the average scores for every 100 bp from the BigWig files were calculated using the rtracklayer package in R (version 1.56.1), and a sliding moving average with a window size of 7 was calculated using the frollmean function from the data.table package (version 1.14.6).

### Reverse transcription-quantitative PCR (RT-qPCR)

RNA purification was performed as described previously^40^. A total of 5×10^7^ cells were disrupted with Mini-Beadbeater-16 in the presence of RNA extraction buffer (20 mM Tris pH 7.5, 300 mM NaCl, 10 mM EDTA, 1% SDS), acid phenol:chloroform:isoamyl alcohol (acid PCI; Sigma, P1944), and glass beads, and RNA was purified by additional acid PCI extraction followed by ethanol precipitation. Residual DNA was removed by 0.05 U/µL RQ1 RNase-Free DNase (Promega, M6101) treatment at 37°C for 30 minutes, followed by acid PCI extraction and ethanol precipitation. Using the purified RNA as a template, complementary DNA (cDNA) was synthesized with High-Capacity cDNA Reverse Transcription Kit (Applied Biosystems, 4368814) and analyzed by qPCR with Fast SYBR Green Master Mix and StepOnePlus Real-Time PCR System. The qPCR primers used in this study are listed in **Supplementary Table 3**.

### Immunofluorescence (IF) microscopy

IF experiments were performed as described previously^41^ with minor modifications to examine the mitotic cell population by visualizing mitotic spindles. A total of 1×10^8^ cells were first fixed with 3% pFA at the culture temperature for 60 minutes and treated with 200 µg/mL Zymolyase-100T (Nacalai Tesque, 07665-55) at 37°C for 20-45 minutes so that more than 90% of the cells were spheroplasted. The nuclei in the spheroplasts were permeabilized by incubation with 1% Triton X-100 at room temperature for 2 minutes. The spheroplasts were next incubated in PEMBAL [100 mM PIPES (Sigma, P1851) pH 6.9, 1 mM ethylene glycol-bis(β-aminoethyl)-N,N,Nʹ,Nʹ-tetraacetic acid (EGTA; Sigma, E3889), 1 mM MgSO_4_ (Sigma, M7506), 1% bovine serum albumin (BSA; Sigma, A7638), 0.1% sodium azide (Sigma, S8032), and 0.1 M L-lysine (Sigma, L5501)] at room temperature for 1 hour, and the anti-tubulin antibody TAT-1^42^ (a gift from Drs.

Gull and Sunter] was added at 1:10 dilution and further incubated at room temperature overnight. After the primary antibody was washed out, Cy3-conjugated anti-Mouse IgG antibody (Jackson ImmunoResearch, 115-165-003) was added at 1:2,000 dilution and incubated at room temperature for 4-6 hours in the dark. The stained cells were mounted on a coverslip with a mounting medium containing 1 µg/mL 4ʹ,6-diamidino-2-phenylindole (DAPI) dilactate (Sigma, D9564) and 100 µg/mL p-phenylenediamine (PPD; Sigma, P6001), and observed under a DeltaVision deconvolution microscope (GE Healthcare).

## DATA AVAILABILITY

All relevant data that support the finding of this study are available from the authors upon request.

## SUPPLEMENTARY INFORMATION

Supplementary Information includes Supplementary Note and Supplementary Table 1-4.

## ACKNOWLEDGMENTS

We would like to thank Drs. Keith Gull and Jack Sunter for anti-tubulin TAT-1 antibody, the University of Oregon Genomics & Cell Characterization Core Facility for next-generation sequencing and microscopic analysis, and the National BioResource Project (NBRP), Japan, for fission yeast strains. We also thank Drs. Jeannie and Eric Selker for scientific and humane guidance, and Ms. Yuko Tsukamoto for technical assistance. This work was supported by the National Institutes of Health/National Institute of General Medical Sciences (R01GM124195 to K.N.), JSPS KAKENHI grant (JP20K23376 to K.N.), Grant for Basic Science Research Projects from The Sumitomo Foundation (K.N.), The Mitsubishi Foundation (K.N.), and Takeda Science Foundation (K.N.).

