## Supplementary for "Local chromatin decompaction shapes mitotic chromosome landscape"

#### EXTENDED DATA FIGURES

##### Extended Data Fig. 1

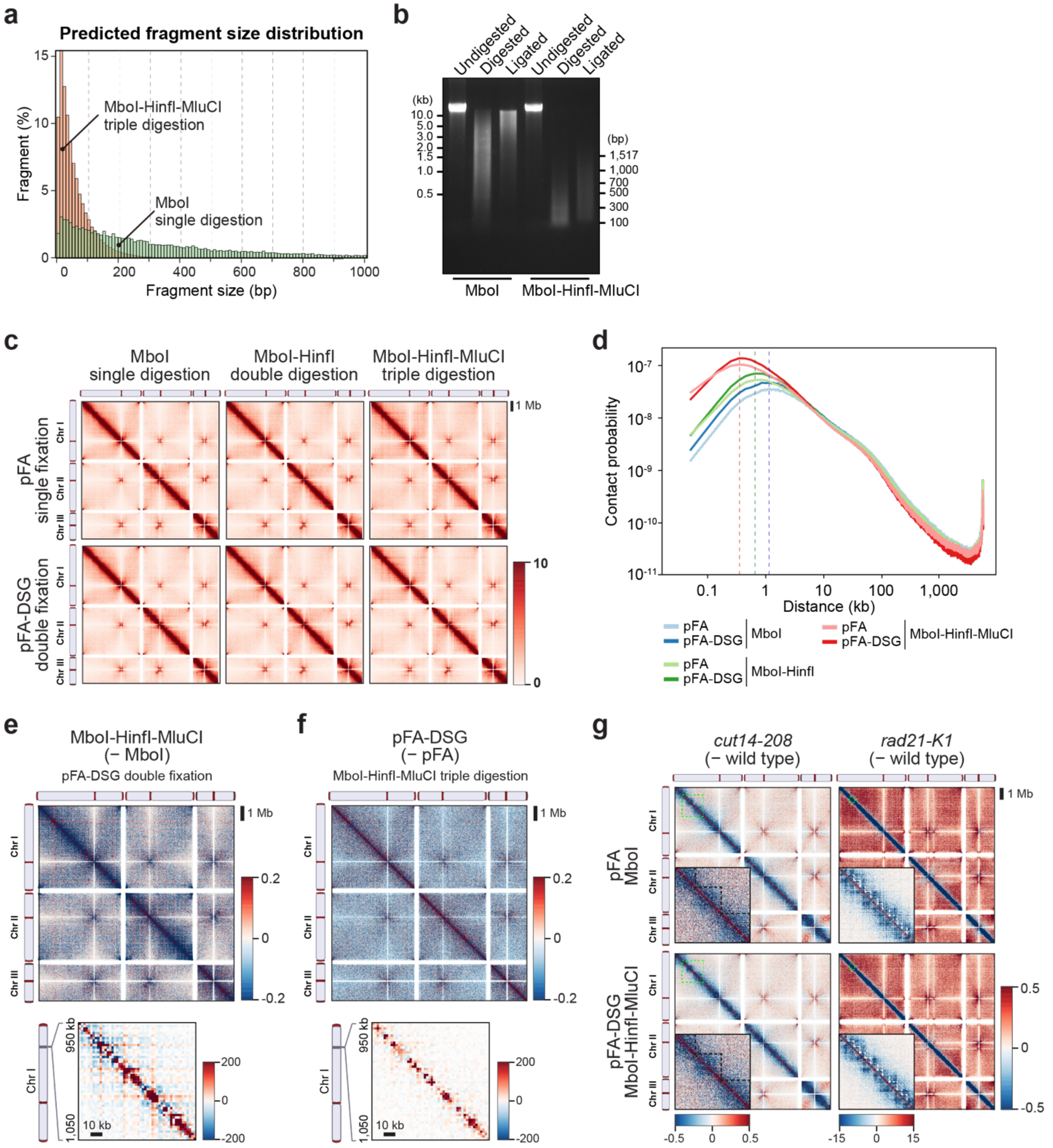


**Extended Data Fig. 1 | Examination of Hi-C conditions.**

**a**, Histograms of the expected size distributions of restriction fragments after MboI single digestion and after MboI-HinfI-MluCI triple digestion of fission yeast genomic DNA.

**b**, DNA size of example Hi-C samples. Hi-C was performed using asynchronously growing fission yeast cells, and DNAs before and after restriction digestion ("Undigested" and "Digested," respectively) and after ligation ("Ligated") were purified and analyzed by agarose gel electrophoresis followed by ethidium bromide staining.

**c**, Hi-C maps of asynchronously growing cells. Hi-C was performed with various restriction digestion and fixation conditions to determine which is the best for detecting short-range contacts.

**d**, Distance-dependent contact probability with various Hi-C conditions used in **Extended Data Fig. 1c**. Dotted lines indicate the peak positions for each restriction digestion condition. Note that the pFA-DSG double fixation condition is always superior to the pFA single fixation in short-range contact detection.

**e**, Differential contacts between the MboI single and MboI-HinfI-MluCI triple restriction digestion conditions. The double fixation condition was commonly used. Below the whole chromosomal map is an enlarged view of the 100-kb region around the *eng1* gene. The increased contact scores along the diagonal (red signals) are indicative of improved detection of short-range contacts with the triple digestion condition.

**f**, Differential contacts between the pFA single and pFA-DSG double fixation conditions. The triple digestion condition was commonly used. Below the whole chromosomal map is an enlarged view of the 100-kb region around the *eng1* gene. The increased contact scores along the diagonal (red signals) are indicative of improved detection of short-range contacts with the double fixation condition.

**g**, Differential contacts upon condensin and cohesin inactivation. Wild-type, *cut14-208* (a temperature-sensitive allele of the condensin subunit gene *cut14*), and *rad21-K1* (a temperature-sensitive allele of the cohesin subunit gene *rad21*) cells growing asynchronously were further cultured at 36°C (a restriction temperature) for 1 h, and Hi-C was performed with either the single fixation, single digestion condition or the double fixation, triple digestion condition. Enlarged views of the 500-kb and 1.9-Mb regions containing the *eng1* gene, as indicated by green dotted line boxes, are shown as insets.

#### Extended Data Fig. 2

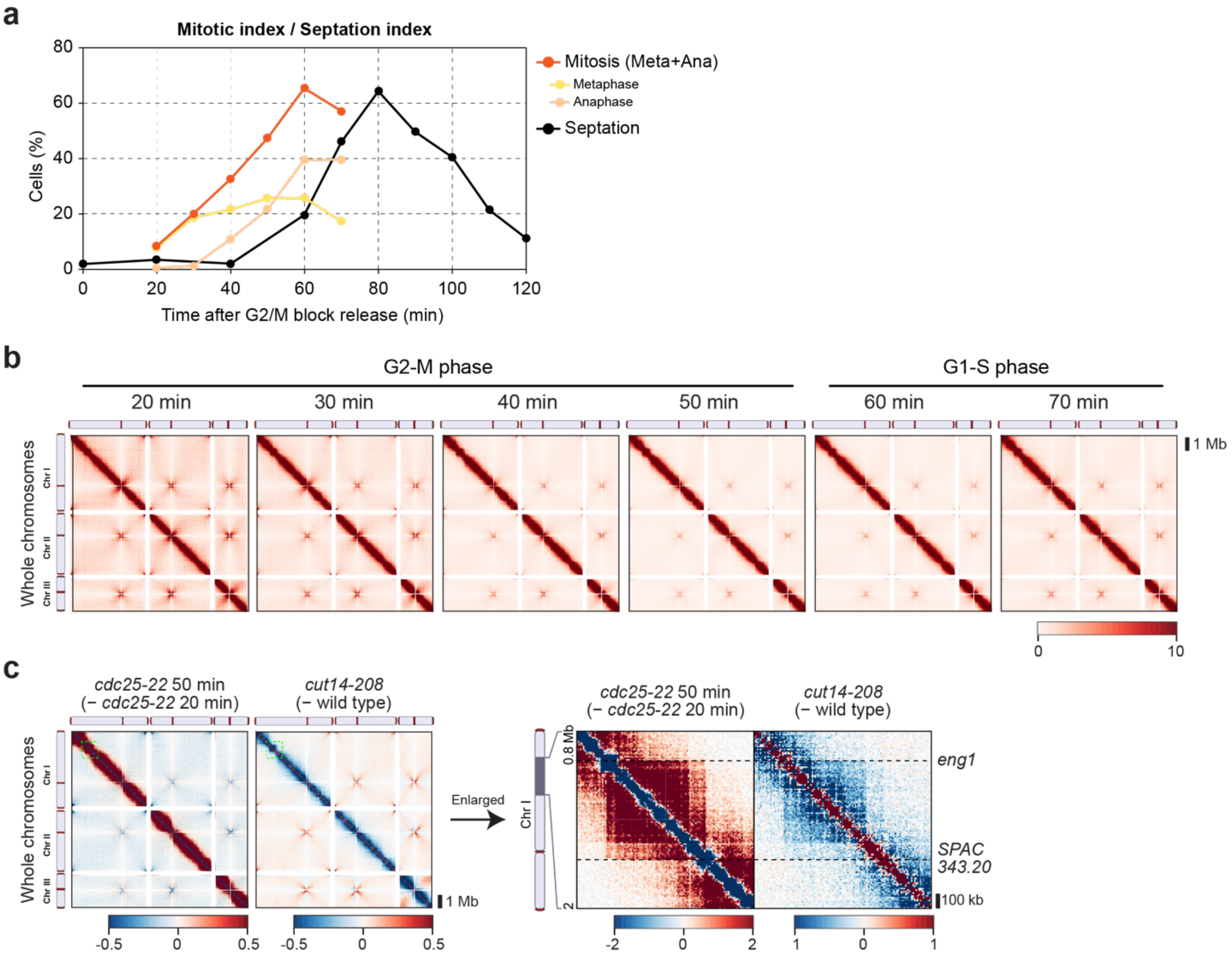


**Extended Data Fig. 2 | Chromosomal reorganization during mitosis after G2/M block release with the *cdc25-22* mutant allele.**

**a**, Mitosis progression after G2/M block release with the *cdc25-22* temperature-sensitive mutant allele. Mitotically synchronized *cdc25-22* cells were fixed every 10 minutes from 20 to 70 minutes after release from the G2/M transition, and immunofluorescence microscopy using the TAT-1 antibody (anti-tubulin antibody) together with DAPI staining was performed to identify metaphase and anaphase cells. The mitotic cell population was determined as a total of metaphase and anaphase cell populations. The septation index, which is known to peak at the S phase, was monitored every 20 minutes after the G2/M block release.

**b**, Time-dependent alteration in chromosome conformation during and after mitosis. The same data as in **Fig. 1a** were used, but Hi-C maps at each time point are shown.

**c**, Condensin-mediated self-associating domains in mitotic cells. Wild-type and *cut14-208* cells growing asynchronously were further cultured at 36°C for 1 h, and Hi-C was performed. Genome-wide Hi-C difference maps between 20- and 50-minute data from the *cdc25-22* block release and between the wild-type and *cut14-208* mutant are shown. Enlarged difference maps of the indicated genomic region (green dotted line boxes in left panels) are shown in the right panels. Highly similar structures between the mitotically formed domains and the lost contacts upon condensin inactivation with the *cut14-208* allele suggest that the mitotic domains are assembled in a condensin-dependent manner[1].

#### Extended Data Fig. 3

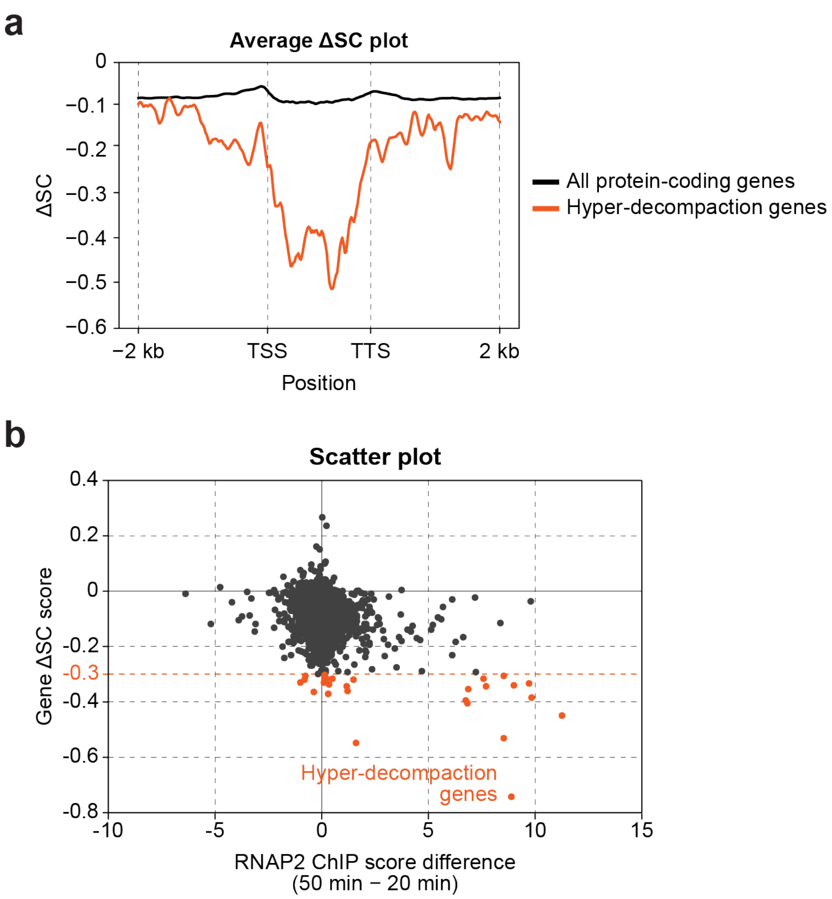


**Extended Data Fig. 3 | Possible involvement of transcription in local decompaction.**

**a**, Hyper-decompaction over the transcribed regions. Average plots of the ΔSC scores over all the protein-coding genes and the hyper-decompaction genes. TSS, transcription start site; TTS, transcription termination site.

**b**, Examination of a possible correlation between mitotic gene activation and local chromatin decompaction levels. The RNAP2 ChIP-seq score difference between 20 and 50 minutes and the gene ΔSC scores at all the protein-coding genes are plotted.

#### Extended Data Fig. 4

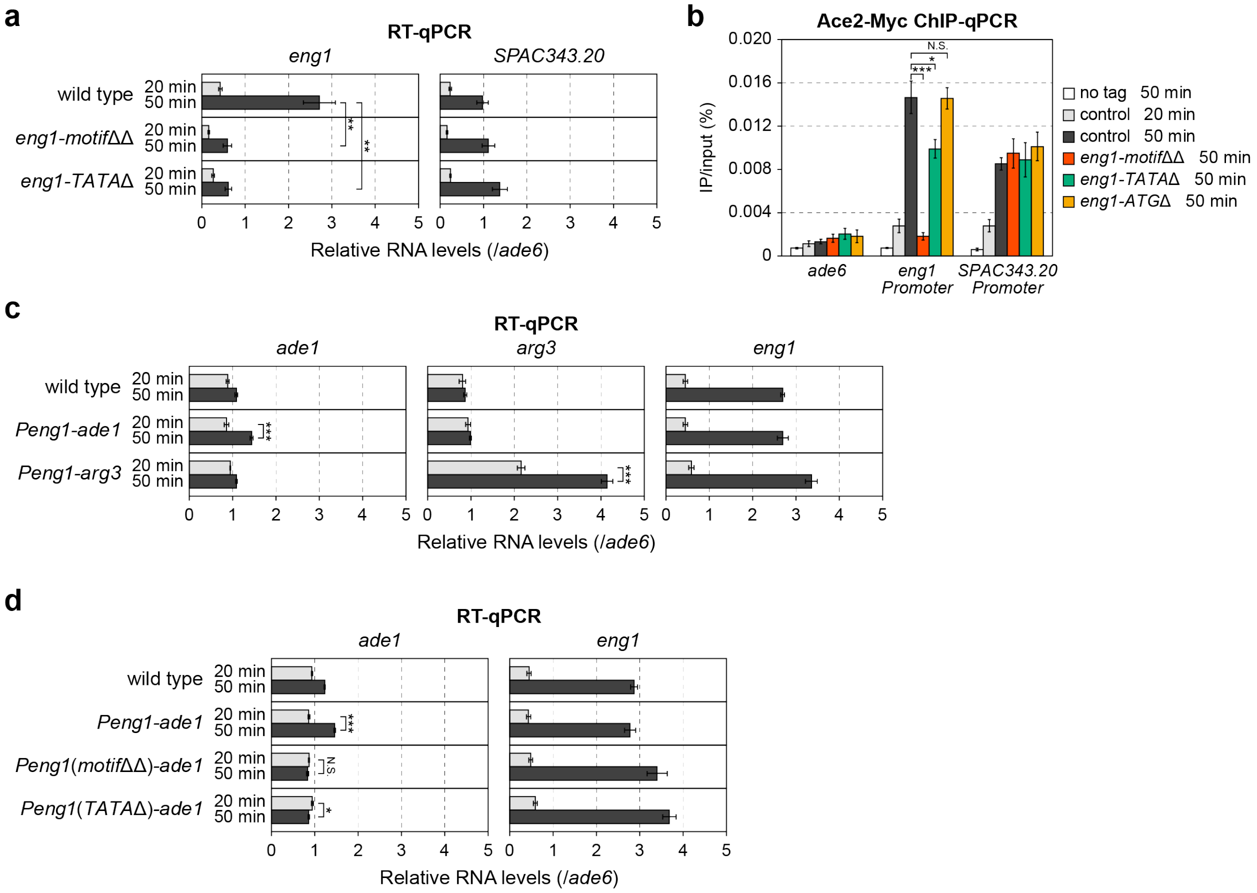


**Extended Data Fig. 4 | Validation of the *eng1* mutant and ectopic *eng1* promoter strains.**

**a**, Gene expression levels in the *eng1* mutants. Mitotically synchronized *cdc25-22* cells in the wild-type, *eng1-motif*ΔΔ, and *eng1-TATA*Δ backgrounds were harvested at 20 and 50 minutes after release from the G2/M transition, and RT-qPCR was performed. The RNA levels of the *SPAC343.20* gene, which undergoes mitotic activation, were also quantified as a control. The RNA levels relative to that of *ade6* were shown, and the error bars represent the standard error of the mean (n=3). Statistical significance was assessed using a two-tailed *t*-test (**, $P < 0.01$).

**b**, Decreased Ace2 binding levels in the *eng1* mutants. Mitotically synchronized *cdc25-22* cells expressing Myc-tagged Ace2 (Ace2-Myc) in the wild-type, *eng1-motif*ΔΔ, and *eng1-TATA*Δ backgrounds were harvested at 50 minutes (also at 20 minutes only for the wild-type control) after release from the G2/M transition, and ChIP-qPCR was performed. Cells with the wild-type *ace2* gene without any tag sequence (no tag) were examined as a negative control. An initial codon-lacking mutant, *eng1-ATG*Δ, was also examined to exclude the possibility that the non-functional *eng1* gene is the cause of the decreased Ace2 binding. DNA recovery after immunoprecipitation relative to input DNA (IP/input) is shown, and the error bars represent the standard error of the mean (n=4). Statistical significance was assessed using a two-tailed *t*-test (N.S., $P > 0.05$; *, $P < 0.05$; ***, $P < 0.001$).

**c,** Forced mitotic activation of the *ade1* and *arg3* genes upon *Peng1* insertion. Mitotically synchronized *cdc25-22* cells in the wild-type, *Peng1-ade1*, and *Peng1-arg3* backgrounds were harvested at 20 and 50 minutes after release from the G2/M transition, and RT-qPCR was performed. The *eng1* RNA level was also quantified as a control gene that undergoes mitotic activation. The RNA levels relative to that of *ade6* were shown, and the error bars represent the standard error of the mean (n=3). Statistical significance was assessed using a two-tailed *t*-test (***, $P < 0.001$).

**d**, Defective mitotic activation of the *ade1* gene when the mutant versions of *Peng1* were used. Mitotically synchronized *cdc25-22* cells in the wild-type, *Peng1-ade1*, *Peng1*(*motif*ΔΔ)-*ade1*, and *Peng1*(*TATA*Δ)-*ade1* backgrounds were harvested at 20 and 50 minutes after release from the G2/M transition, and RT-qPCR was performed. The *eng1* RNA level was also quantified as a control gene that undergoes mitotic activation. The RNA levels relative to that of *ade6* were shown, and the error bars represent the standard error of the mean (n=3). Statistical significance was assessed using a two-tailed *t*-test (N.S., $P > 0.05$; *, $P < 0.05$; ***, $P < 0.001$).

#### Extended Data Fig. 5

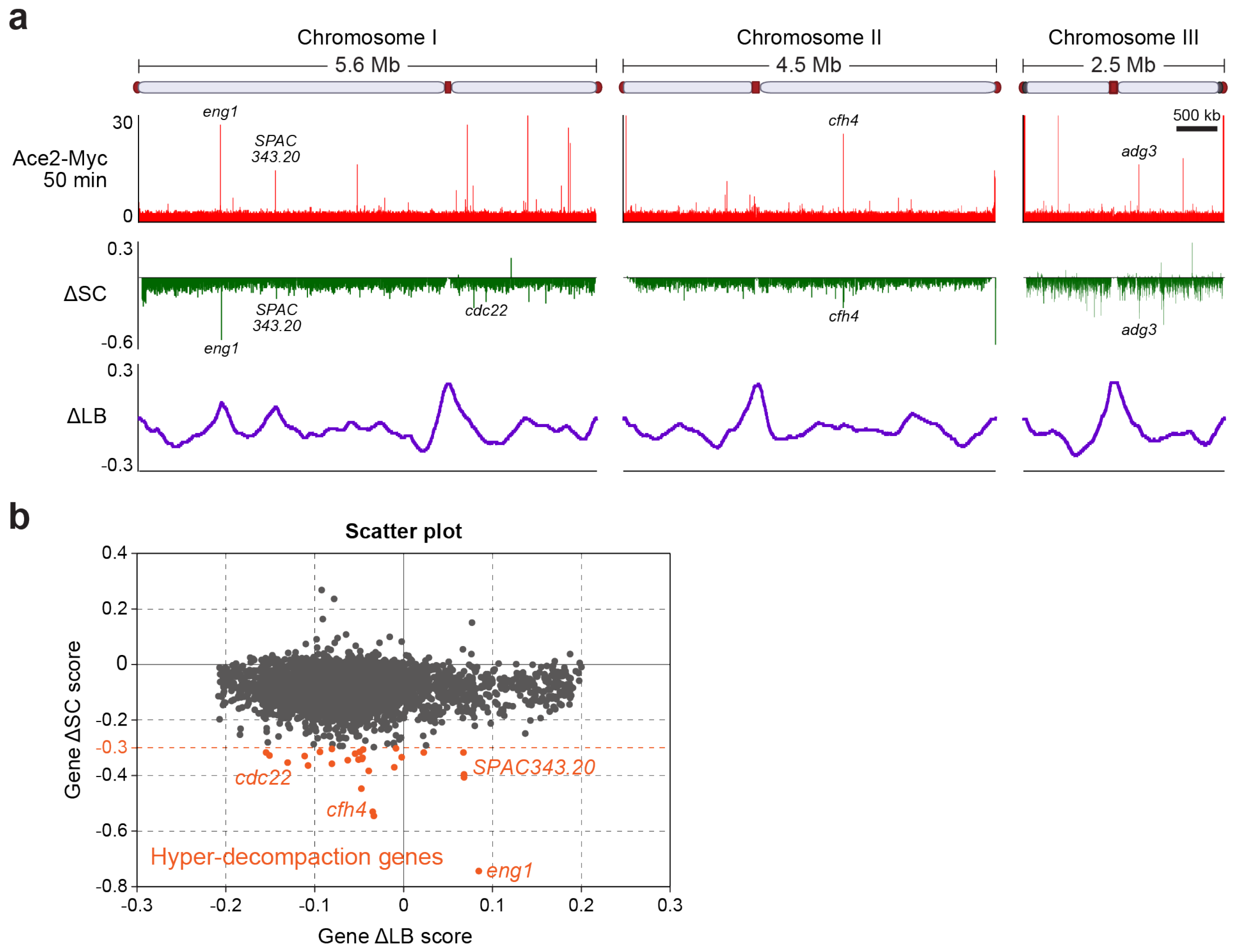


**Extended Data Fig. 5 | Examination of possible correlation between local decompaction and boundary formation levels.**

**a**, A comparison of the genome-wide profiles of the ΔSC and ΔLB scores. The genome-wide Ace2-Myc distribution is also shown to test a possible coincidence among the ΔSC, ΔLB, and Ace2 peaks.

**b,** Examination of a possible correlation between compaction and boundary levels. The gene ΔSC and gene ΔLB scores at all the protein-coding genes are plotted. Gene ΔLB scores did not show an obvious correlation with gene ΔSC scores, which is likely due to the significantly broader peaks of ΔLB scores and the chromatin context dependency of boundary formation, calling for careful experimental settings to examine the possible connection between ΔLB and ΔSC scores.

#### Extended Data Fig. 6

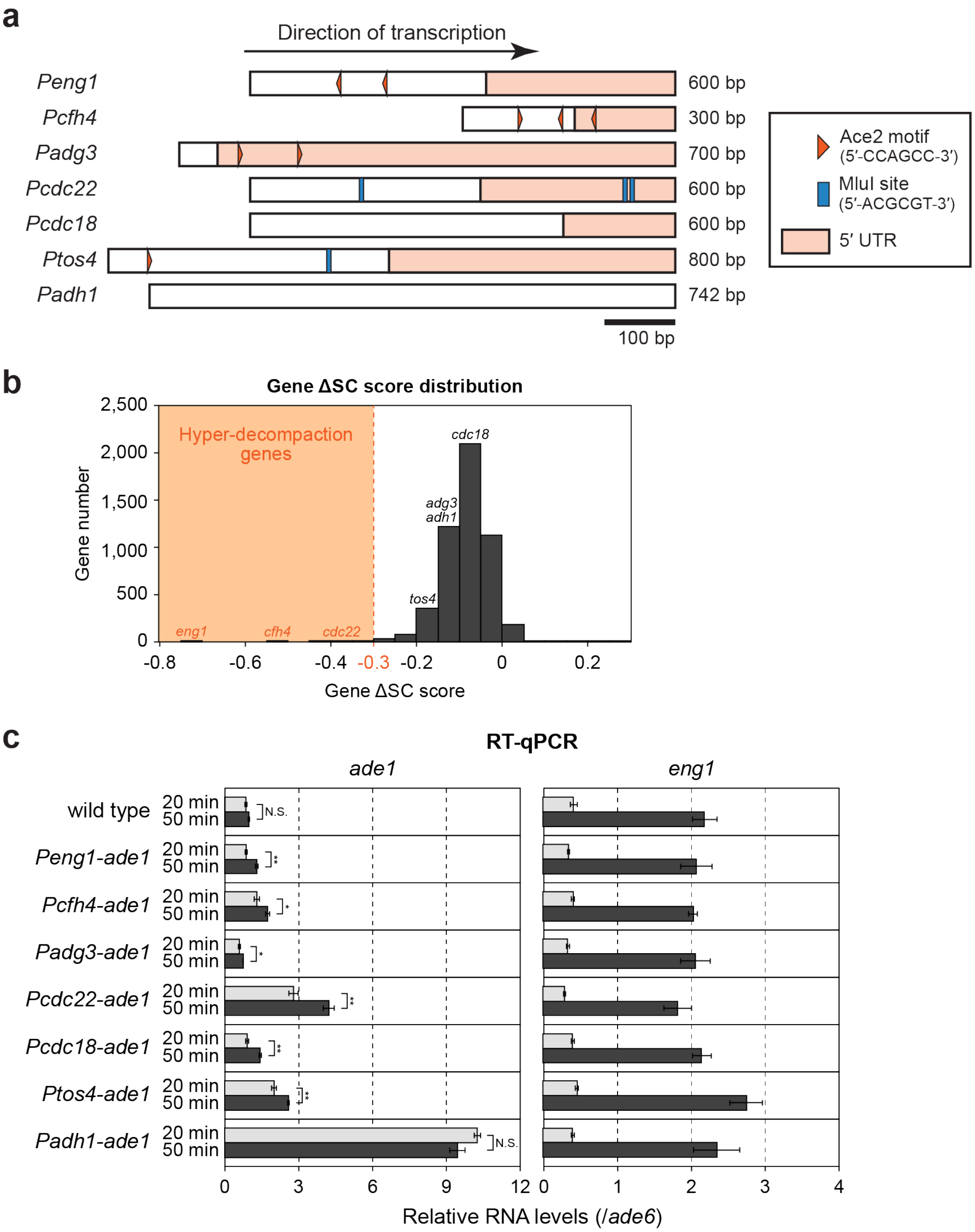


**Extended Data Fig. 6 | Validation of the ectopic promoter strains.**

**a**, The promoter sequences used for the ectopic promoter strains in this study. The positions of Ace2 binding motifs (5′-CCAGCC-3′) and the MluI restriction sites (5′-ACGCGT-3′) that can be targeted by the MBF transcription factor are indicated by the red arrowheads and blue boxes, respectively. The orange boxes indicate 5′ untranslated regions (UTR).

**b**, Histograms of the gene ΔSC score distribution. The names of genes whose promoters were used for the ectopic promoter strains are indicated over the bin to which they belong.

**c**, Forced mitotic activation of the *ade1* gene in the ectopic promoter strains. Mitotically synchronized *cdc25-22* cells that harbor the *ade1* gene with or without the ectopic promoter insertion were harvested at 20 and 50 minutes after release from the G2/M transition, and RT-qPCR was performed. The *eng1* RNA level was also quantified as a control gene that undergoes mitotic activation. The RNA levels relative to that of *ade6* were shown, and the error bars represent the standard error of the mean (n=3). Statistical significance was assessed using a two-tailed *t*-test (N.S., $P > 0.05$; *, $P < 0.05$; **, $P < 0.01$).

#### Extended Data Fig. 7

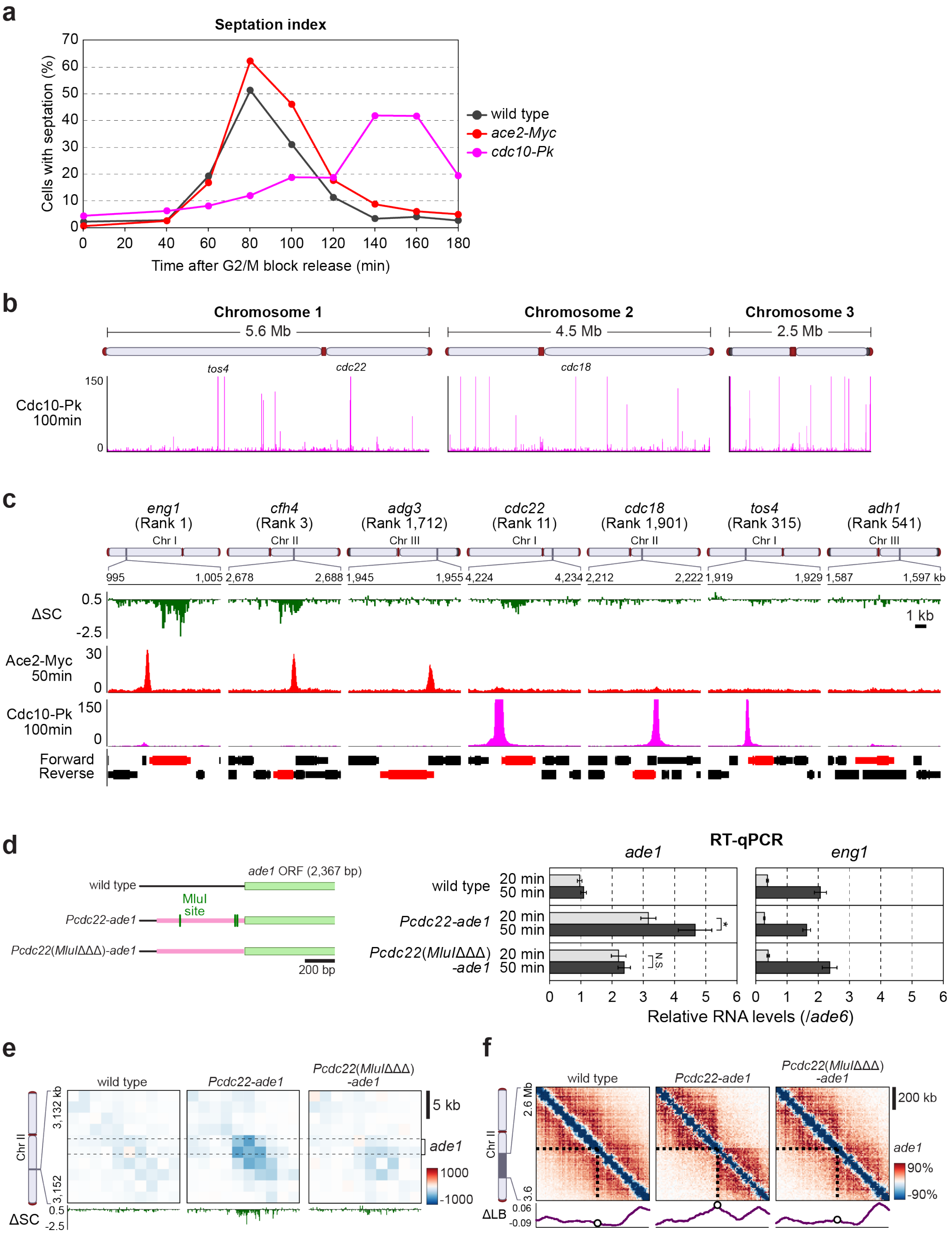


**Extended Data Fig. 7 | Potential of the MBF transcription factor in local decompaction and**

**boundary formation.**

**a**, Delayed cell cycle progression of cells expressing epitope-tagged Cdc10 proteins. The septation index of *cdc25-22* cells in the wild-type, *ace2-Myc*, and *cdc10-Pk* backgrounds after G2/M block release was monitored.

**b**, A genome-wide binding profile of Cdc10, a subunit of the MBF transcription factor complex, during mitosis. Mitotically synchronized *cdc25-22* cells expressing Cdc10-Pk from the endogenous promoter were fixed at 100 minutes after release from the G2/M transition, and ChIP-seq was performed using anti-Pk antibody. Cells at 100-minute time point instead of the usual 50-minute were examined because of the delayed cell cycle progression after the block release (**Extended Data Fig. 7a**).

**c,** Binding of Ace2 and Cdc10 to genes whose promoters were used for the ectopic promoter strains. The Ace2-Myc and Cdc10-Pk ChIP-seq plots at each gene locus were shown. Above the ChIP-seq plots is the ΔSC score plot for comparison.

**d**, Defective mitotic activation of the *ade1* gene when the mutant version of *Pcdc22* was used. Mitotically synchronized *cdc25-22* cells in the wild-type, *Pcdc22-ade1*, and *Pcdc22*(*MluI*ΔΔΔ)-*ade1* backgrounds were harvested at 20 and 50 minutes after release from the G2/M transition, and RT-qPCR was performed. The *eng1* RNA level was also quantified as a control gene that undergoes mitotic activation. The RNA levels relative to that of *ade6* were shown, and the error bars represent the standard error of the mean (n=5). Statistical significance was assessed using a two-tailed *t*-test (N.S., $P > 0.05$; *, $P < 0.05$).

**e**, Impaired local decompaction upon insertion of the mutant *Pcdc22*. Differential contacts between 20 and 50 minutes in each strain are shown for the 20-kb region containing the *ade1* gene, and below each Hi-C difference map is its ΔSC score plot.

**f**, Defective formation of long-range contact boundary upon insertion of the mutant *Pcdc22*. Differential contacts between 20 and 50 minutes in each strain are shown for the indicated 1-Mb region of chromosome II, and below each Hi-C map is its ΔLB score plot.

#### Extended Data Fig. 8

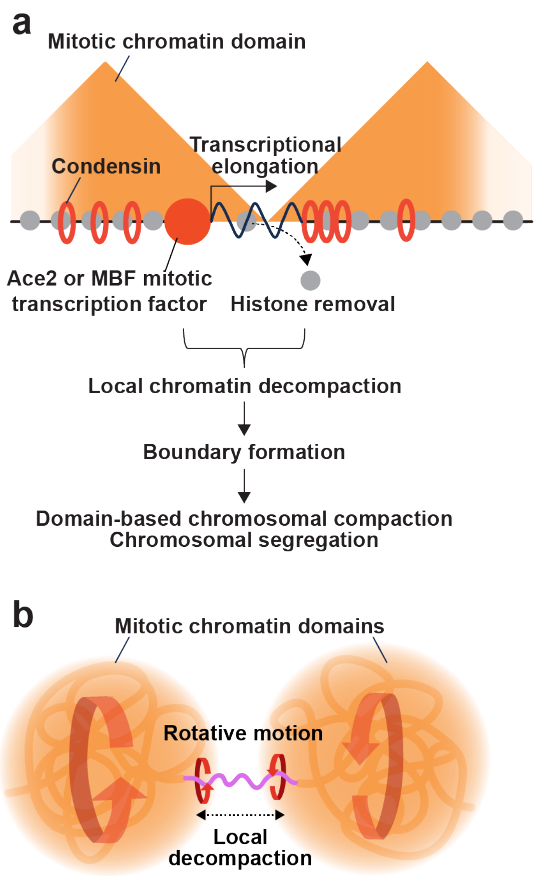


**Extended Data Fig. 8 | Schematic models.**

**a**, A model to connect local chromatin decompaction to large-scale chromosomal reorganization during mitosis; 1) Transcriptional elongation of Ace2- or MBF-driven mitotically activated genes is required for local chromatin decompaction; 2) Local chromatin decompaction is coupled with histone (nucleosomal) removal. 3) Mitotically activated genes form boundaries of mitotic chromatin domains that serve as chromosomal compaction units; 4) Domain-based chromosomal compaction allows for faithful chromosomal segregation.

**b**, A model to explain why local chromatin decompaction coupled with transcriptional elongation establishes domain boundaries. Since Ace2- or MBF-driven transcription and its elongation distance are essential parameters, we hypothesize that local chromatin decompaction not only physically separates two adjoining domains but also prevents inter-domain contacts by a rotative motion during transcriptional elongation.

#### SUPPLEMENTARY INFORMATION

##### Supplementary Note | Exploration of Hi-C conditions for better detection of fine-scale DNA contacts.

To fragment genomic DNA into smaller pieces, we selected three enzymes, MboI (5′-GATC-3′), HinfI (5′-GANTC-3′), and MluCI (5′-AATT-3′), because these enzymes are highly active in 1× NEBuffer 2 (New England Biolabs, B7002S) and also because the resultant protruding ends after digestion are compatible with DNA-end labeling with biotin-conjugated dATP. Fission yeast genomic DNA has 32,831, 39,079, and 142,215 sites that are recognized by MboI, HinfI, and MluCI, respectively, and expected sizes of DNA fragments with MboI single digestion and MboI-HinfI-MluCI triple digestion are 383 and 59 bp on average, respectively (**Extended Data Fig. 1a**). As expected, the triple digestion resulted in much smaller DNA fragments compared to the single digestion (**Extended Data Fig. 1b**).

We first conducted in situ Hi-C experiments with MboI single, MboI-HinfI double[1], and MboI-HinfI-MluCI triple digestions. In addition, we also examined two chromatin fixation conditions in this study, one with paraformaldehyde (pFA), which is commonly used for Hi-C, and the other with disuccinimidyl glutarate (DSG) in addition to pFA, because a previously reported Hi-C derivative with nucleosome-resolution, termed Micro-C, cannot efficiently capture long-range contacts without DSG fixation[2,3]. In our experiments, however, long-range contacts, including the inter-centromere and inter-telomere contacts, were clearly detected with all these fixation and digestion conditions (**Extended Data Fig. 1c**), although the triple digestion slightly inefficiently captured long-range contacts beyond 100 kb (**Extended Data Fig. 1d**). As we expected, the MboI-HinfI-MluCI triple digestion was able to detect short-range contacts more efficiently than the MboI single and MboI-HinfI double digestions, as evidenced by the higher contact probability in short-range contacts (**Extended Data Fig. 1d,e**). Moreover, we found that the pFA-DSG double fixation was superior to the pFA single fixation in detecting short-range contacts less (**Extended Data Fig. 1d,f**). We therefore decided to use the MboI-HinfI-MluCI triple digestion, pFA-DSG double fixation condition for further analyses in this study.

#### Supplementary Table 1 | Hyper-decompaction gene list.

| systematic name | gene name | chr | start | end | size (bp) | ΔSC rank | ΔSC score | M up-regulation | Ace2 binding | Cdc10 binding | Note |
|---|---|---|---|---|---|---|---|---|---|---|---|
| SPAC821.09 | *eng1* | I | 998,632 | 1,002,361 | 3,729 | 1 | -0.744487 | Yes | Yes | | |
| SPBC3E7.13c | *syf2* | II | 2,683,836 | 2,684,840 | 1,004 | 2 | -0.547057 | | | | Adjacent to cfh4 |
| SPBC3E7.12c | *cfh4* | II | 2,682,000 | 2,683,804 | 1,804 | 3 | -0.531427 | Yes | Yes | | |
| SPCC622.09 | *htb1* | III | 1,412,359 | 1,413,677 | 1,318 | 4 | -0.449082 | Yes | Yes | Yes | |
| SPAC343.12 | *rds1* | I | 1,667,739 | 1,669,666 | 1,927 | 5 | -0.406965 | Yes | | | Adjacent to SPAC343.20 |
| SPAC343.20 | | I | 1,667,955 | 1,669,964 | 2,009 | 6 | -0.396329 | Yes | Yes | | |
| SPAC1B3.16c | *vht1* | I | 4,958,698 | 4,960,910 | 2,212 | 7 | -0.383753 | Yes | | | |
| SPAC977.09c | *plb4* | I | 45,875 | 48,160 | 2,285 | 8 | -0.370317 | | | | |
| SPCC191.02c | *acs1* | III | 1,708,234 | 1,711,217 | 2,983 | 9 | -0.364156 | | | | |
| SPAC11E3.13c | *gas5* | I | 5,306,742 | 5,309,045 | 2,303 | 10 | -0.359589 | | | | |
| SPAC1F7.05 | *cdc22* | I | 4,226,970 | 4,229,989 | 3,019 | 11 | -0.353926 | Yes | | Yes | |
| SPAPJ760.03c | *adg1* | I | 5,265,435 | 5,266,485 | 1,050 | 12 | -0.344962 | Yes | Yes | | |
| SPBC2G5.05 | *tkt1* | II | 2,586,380 | 2,588,834 | 2,454 | 13 | -0.344581 | | | | |
| SPAC19G12.16c | *adg2* | I | 4,078,389 | 4,081,156 | 2,767 | 14 | -0.342026 | Yes | Yes | | |
| SPAC31G5.11 | *pac2* | I | 3,006,138 | 3,006,845 | 707 | 15 | -0.337284 | | | | |
| SPAPYUG7.03c | *mid2* | I | 4,748,861 | 4,751,453 | 2,592 | 16 | -0.334151 | Yes | Yes | | |
| SPCC576.11 | *rpl15* | III | 2,099,846 | 2,100,555 | 709 | 17 | -0.33102 | | | | |
| SPBC14F5.05c | *sam1* | II | 4,162,069 | 4,163,858 | 1,789 | 18 | -0.329045 | | | | |
| SPAC22H10.13 | *zym1* | I | 2,381,666 | 2,382,200 | 534 | 19 | -0.321538 | | | | |
| SPAC343.21 | | I | 1,666,787 | 1,667,735 | 948 | 20 | -0.31877 | | | | Adjacent to SPAC343.20 |
| SPBC32H8.03 | *bem46* | II | 1,457,486 | 1,458,849 | 1,363 | 21 | -0.318584 | | | | |
| SPCC1020.06c | *tal1* | III | 773,366 | 774,834 | 1,468 | 22 | -0.317933 | | | | |
| SPAC14C4.09 | *agn1* | I | 5,244,037 | 5,245,966 | 1,929 | 23 | -0.315907 | Yes | Yes | | |
| SPCC1442.08c | *cox12* | III | 1,781,404 | 1,782,074 | 670 | 24 | -0.315609 | | | | |
| SPCC622.08c | *hta1* | III | 1,411,144 | 1,412,181 | 1,037 | 25 | -0.307536 | Yes | Yes | Yes | |
| SPAC3C7.14c | *obr1* | I | 2,094,110 | 2,094,984 | 874 | 26 | -0.30595 | | | | |
| SPAC977.03 | | I | 33,835 | 34,272 | 437 | 27 | -0.302032 | | | | |

M up-regulation (mitotic gene up-regulation), “Yes” if the RNAP2 ChIP-seq score difference between 20 and 50 minutes of the gene of interest is more than 2 (**Extended Data Fig. 3b**). Ace2 binding / Cdc10 binding, “Yes” if Ace2-Myc / Cdc10-Pk ChIP-seq peaks near the 5′ UTR region of the gene of interest (**Fig. 2a** and **Extended Data Fig. 7b**).

**Supplementary Table 2 | *S. pombe* strains used in this study.**

| Figure<br>Strains | Genotype |
|---|---|
| **Fig. 1a-c,e,g** | |
| SPSTn249 | *mat1M ade6-M216 his2 leu1-32 ura4 cdc25-22* |
| **Fig. 1h** | |
| SPSTn249 | *mat1M ade6-M216 his2 leu1-32 ura4 cdc25-22* |
| SPSTn494 | *mat1M ade6-M216 his2 leu1-32 ura4 cdc25-22 cut14-12Pk-kanMX6* |
| **Fig. 2a** | |
| SPSTn462 | *mat1M ade6-M216 his2 leu1-32 ura4-D18 cdc25-22 ace2-13Myc-kanMX6* |
| **Fig. 2b** | |
| SPSTn249 | *mat1M ade6-M216 his2 leu1-32 ura4 cdc25-22* |
| SPSTn462 | *mat1M ade6-M216 his2 leu1-32 ura4-D18 cdc25-22 ace2-13Myc-kanMX6* |
| **Fig. 2c,f** | |
| SPSTn249 | *mat1M ade6-M216 his2 leu1-32 ura4 cdc25-22* |
| SPSTn479 | *mat1M ade6-M210 his2 leu1-32 ura4 cdc25-22 eng1-TATA*Δ |
| SPSTn750 | *mat1M ade6-M216 his2 leu1-32 ura4 cdc25-22 eng1-motif*ΔΔ |
| **Fig. 2d,g** | |
| SPSTn249 | *mat1M ade6-M216 his2 leu1-32 ura4 cdc25-22* |
| SPSTn477 | *mat1M ade6-M216 his2 leu1-32 ura4 cdc25-22 natMX6-Peng1-ade1* |
| **Fig. 2e,h** | |
| SPSTn249 | *mat1M ade6-M216 his2 leu1-32 ura4 cdc25-22* |
| SPSTn475 | *mat1M ade6-M216 his2 leu1-32 ura4 cdc25-22 natMX6-Peng1-arg3* |

**Fig. 3a,b**

SPSTn249 *mat1M ade6-M216 his2 leu1-32 ura4 cdc25-22*

SPSTn477 *mat1M ade6-M216 his2 leu1-32 ura4 cdc25-22 natMX6-Peng1-ade1*

SPSTn787 *mat1M ade6-M216 his2 leu1-32 ura4 cdc25-22 natMX6-Peng1*(*TATA*Δ)*-ade1*

SPSTn788 *mat1M ade6-M216 his2 leu1-32 ura4 cdc25-22 natMX6-Peng1*(*motif*ΔΔ)*-ade1*

**Fig. 3d**

SPSTn249 *mat1M ade6-M216 his2 leu1-32 ura4 cdc25-22*

**Fig. 4a**

SPSTn249 *mat1M ade6-M216 his2 leu1-32 ura4 cdc25-22*

SPSTn462 *mat1M ade6-M216 his2 leu1-32 ura4-D18 cdc25-22 ace2-13Myc-kanMX6*

**Fig. 4b,c**

SPSTn249 *mat1M ade6-M216 his2 leu1-32 ura4 cdc25-22*

SPSTn477 *mat1M ade6-M216 his2 leu1-32 ura4 cdc25-22 natMX6-Peng1-ade1*

SPSTn478 *mat1M ade6-M210 his2 leu1-32 ura4 cdc25-22 natMX6-Padh1-ade1*

SPSTn736 *mat1M ade6-M216 his2 leu1-32 ura4 cdc25-22 natMX6-Ptos4-ade1*

SPSTn737 *mat1M ade6-M216 his2 leu1-32 ura4 cdc25-22 natMX6-Pcdc22-ade1*

SPSTn739 *mat1M ade6-M216 his2 leu1-32 ura4 cdc25-22 natMX6-Pcdc18-ade1*

SPSTn740 *mat1M ade6-M216 his2 leu1-32 ura4 cdc25-22 natMX6-Pcfh4-ade1*

SPSTn741 *mat1M ade6-M216 his2 leu1-32 ura4 cdc25-22 natMX6-Padg3-ade1*

**Fig. 4d**

| | |
|---|---|
| SPSTn249 | *mat1M ade6-M216 his2 leu1-32 ura4 cdc25-22* |
| SPSTn477 | *mat1M ade6-M216 his2 leu1-32 ura4 cdc25-22 natMX6-Peng1-ade1* |
| SPSTn478 | *mat1M ade6-M210 his2 leu1-32 ura4 cdc25-22 natMX6-Padh1-ade1* |
| SPSTn736 | *mat1M ade6-M216 his2 leu1-32 ura4 cdc25-22 natMX6-Ptos4-ade1* |
| SPSTn737 | *mat1M ade6-M216 his2 leu1-32 ura4 cdc25-22 natMX6-Pcdc22-ade1* |
| SPSTn739 | *mat1M ade6-M216 his2 leu1-32 ura4 cdc25-22 natMX6-Pcdc18-ade1* |
| SPSTn740 | *mat1M ade6-M216 his2 leu1-32 ura4 cdc25-22 natMX6-Pcfh4-ade1* |
| SPSTn741 | *mat1M ade6-M216 his2 leu1-32 ura4 cdc25-22 natMX6-Padg3-ade1* |
| SPSTn787 | *mat1M ade6-M216 his2 leu1-32 ura4 cdc25-22 natMX6-Peng1*(*TATA*Δ)*-ade1* |
| SPSTn788 | *mat1M ade6-M216 his2 leu1-32 ura4 cdc25-22 natMX6-Peng1*(*motif*ΔΔ)*-ade1* |
| SPSTn835 | *mat1M ade6-M216 his2 leu1-32 ura4 cdc25-22 natMX6-Pcdc22*(*MluI*ΔΔΔ)*-ade1* |

**Fig. 5a-c**

| | |
|---|---|
| SPSTn249 | *mat1M ade6-M216 his2 leu1-32 ura4 cdc25-22* |
| SPSTn1153 | *mat1M ade6-M216 his2 leu1-32 ura4 cdc25-22 eng1*:$T_{CYC1}$(*1*) |
| SPSTn1155 | *mat1M ade6-M216 his2 leu1-32 ura4 cdc25-22 eng1*:$T_{CYC1}$(*1086*) |
| SPSTn1157 | *mat1M ade6-M216 his2 leu1-32 ura4 cdc25-22 eng1*:$T_{CYC1}$(*2247*) |
| SPSTn1175 | *mat1M ade6-M216 his2 leu1-32 ura4 cdc25-22 eng1*:$T_{CYC1}$(*3369*) |

**Fig. 5d**

SPSTn249 *mat1M ade6-M216 his2 leu1-32 ura4 cdc25-22*

SPSTn479 *mat1M ade6-M210 his2 leu1-32 ura4 cdc25-22 eng1-TATAΔ*

SPSTn750 *mat1M ade6-M216 his2 leu1-32 ura4 cdc25-22 eng1-motifΔΔ*

SPSTn1153 *mat1M ade6-M216 his2 leu1-32 ura4 cdc25-22 eng1*:$T_{CYC1}$(*1*)

SPSTn1155 *mat1M ade6-M216 his2 leu1-32 ura4 cdc25-22 eng1*:$T_{CYC1}$(*1086*)

SPSTn1157 *mat1M ade6-M216 his2 leu1-32 ura4 cdc25-22 eng1*:$T_{CYC1}$(*2247*)

SPSTn1175 *mat1M ade6-M216 his2 leu1-32 ura4 cdc25-22 eng1*:$T_{CYC1}$(*3369*)

**Fig. 6a-c**

SPSTn249 *mat1M ade6-M216 his2 leu1-32 ura4 cdc25-22*

**Fig. 6d**

SPSTn249 *mat1M ade6-M216 his2 leu1-32 ura4 cdc25-22*

SPSTn479 *mat1M ade6-M210 his2 leu1-32 ura4 cdc25-22 eng1-TATAΔ*

SPSTn750 *mat1M ade6-M216 his2 leu1-32 ura4 cdc25-22 eng1-motifΔΔ*

**Extended Data Fig. 1b-f**

SPSTn1 $h^{+N}$ *ade6-M210 leu1-32 ura4-DS/E*

**Extended Data Fig. 1g**

SPSTn2 $h^{-}$ *leu1-32 cut14-208*

SPSTn30 $h^{-}$

SPSTn152 $h^{-}$ *leu1 ura4 rad21*::*rad21-K1*(*ura4*$^{+}$)

**Extended Data Fig. 2a,b**

SPSTn249 *mat1M ade6-M216 his2 leu1-32 ura4 cdc25-22*

**Extended Data Fig. 2c**

SPSTn2 *h⁻ leu1-32 cut14-208*

SPSTn30 *h⁻*

SPSTn249 *mat1M ade6-M216 his2 leu1-32 ura4 cdc25-22*

**Extended Data Fig. 3a,b**

SPSTn249 *mat1M ade6-M216 his2 leu1-32 ura4 cdc25-22*

**Extended Data Fig. 4a**

SPSTn249 *mat1M ade6-M216 his2 leu1-32 ura4 cdc25-22*

SPSTn479 *mat1M ade6-M210 his2 leu1-32 ura4 cdc25-22 eng1-TATAΔ*

SPSTn750 *mat1M ade6-M216 his2 leu1-32 ura4 cdc25-22 eng1-motifΔΔ*

**Extended Data Fig. 4b**

SPSTn249 *mat1M ade6-M216 his2 leu1-32 ura4 cdc25-22*

SPSTn462 *mat1M ade6-M216 his2 leu1-32 ura4-D18 cdc25-22 ace2-13Myc-kanMX6*

SPSTn486 *mat1M ade6-M216 his2 leu1-32 ura4-D18 cdc25-22 eng1-TATAΔ ace2-13Myc-kanMX6*

SPSTn827 *mat1M ade6-M216 his2 leu1-32 ura4-D18 cdc25-22 eng1-motifΔΔ ace2-13Myc-kanMX6*

SPSTn829 *mat1M ade6-M216 his2 leu1-32 ura4-D18 cdc25-22 eng1-ATGΔ ace2-13Myc-kanMX6*

**Extended Data Fig. 4c**

SPSTn249 *mat1M ade6-M216 his2 leu1-32 ura4 cdc25-22*

SPSTn475 *mat1M ade6-M216 his2 leu1-32 ura4 cdc25-22 natMX6-Peng1-arg3*

SPSTn477 *mat1M ade6-M216 his2 leu1-32 ura4 cdc25-22 natMX6-Peng1-ade1*

**Extended Data Fig. 4d**

SPSTn249 *mat1M ade6-M216 his2 leu1-32 ura4 cdc25-22*

SPSTn477 *mat1M ade6-M216 his2 leu1-32 ura4 cdc25-22 natMX6-Peng1-ade1*

SPSTn787 *mat1M ade6-M216 his2 leu1-32 ura4 cdc25-22 natMX6-Peng1*(*TATA*Δ)*-ade1*

SPSTn788 *mat1M ade6-M216 his2 leu1-32 ura4 cdc25-22 natMX6-Peng1*(*motif*ΔΔ)*-ade1*

**Extended Data Fig. 5a**

SPSTn249 *mat1M ade6-M216 his2 leu1-32 ura4 cdc25-22*

SPSTn462 *mat1M ade6-M216 his2 leu1-32 ura4-D18 cdc25-22 ace2-13Myc-kanMX6*

**Extended Data Fig. 5b**

SPSTn249 *mat1M ade6-M216 his2 leu1-32 ura4 cdc25-22*

**Extended Data Fig. 6b**

SPSTn249 *mat1M ade6-M216 his2 leu1-32 ura4 cdc25-22*

**Extended Data Fig. 6c**

SPSTn249 *mat1M ade6-M216 his2 leu1-32 ura4 cdc25-22*

SPSTn477 *mat1M ade6-M216 his2 leu1-32 ura4 cdc25-22 natMX6-Peng1-ade1*

SPSTn478 *mat1M ade6-M210 his2 leu1-32 ura4 cdc25-22 natMX6-Padh1-ade1*

| | |
|---|---|
| SPSTn736 | *mat1M ade6-M216 his2 leu1-32 ura4 cdc25-22 natMX6-Ptos4-ade1* |
| SPSTn737 | *mat1M ade6-M216 his2 leu1-32 ura4 cdc25-22 natMX6-Pcdc22-ade1* |
| SPSTn739 | *mat1M ade6-M216 his2 leu1-32 ura4 cdc25-22 natMX6-Pcdc18-ade1* |
| SPSTn740 | *mat1M ade6-M216 his2 leu1-32 ura4 cdc25-22 natMX6-Pcfh4-ade1* |
| SPSTn741 | *mat1M ade6-M216 his2 leu1-32 ura4 cdc25-22 natMX6-Padg3-ade1* |

**Extended Data Fig. 7a,c**

| | |
|---|---|
| SPSTn249 | *mat1M ade6-M216 his2 leu1-32 ura4 cdc25-22* |
| SPSTn462 | *mat1M ade6-M216 his2 leu1-32 ura4-D18 cdc25-22 ace2-13Myc-kanMX6* |
| SPSTn1016 | *mat1M ade6-M216 his2 leu1-32 ura4 cdc25-22 cdc10-4Pk-kanMX6* |

**Extended Data Fig. 7b**

| | |
|---|---|
| SPSTn1016 | *mat1M ade6-M216 his2 leu1-32 ura4 cdc25-22 cdc10-4Pk-kanMX6* |

**Extended Data Fig. 7d-f**

| | |
|---|---|
| SPSTn249 | *mat1M ade6-M216 his2 leu1-32 ura4 cdc25-22* |
| SPSTn737 | *mat1M ade6-M216 his2 leu1-32 ura4 cdc25-22 natMX6-Pcdc22-ade1* |
| SPSTn835 | *mat1M ade6-M216 his2 leu1-32 ura4 cdc25-22 natMX6-Pcdc22*(*MluI*ΔΔΔ)-*ade1* |

The original names of the SPSTn2, SPSTn30, and SPSTn152 strains are FY8027, FY7507, and FY11001, respectively, which were provided by the National BioResource Project (NBRP), Japan.

**Supplementary Table 3 | Primers used in this study.**

---

**Primers for strain constructions**

---

***eng1-motif*ΔΔ construction**

STn407 5′-CCAAACAATATGTTGCCTGTAC-3′

STn537 5′-GGAAATCTGGATGTCTTAACAGCTTTTTAAGAAAAACTGTTTATACATAAGCA-3′

STn538 5′-CACATTTGTCCTCTTTATGGTAAATATACATGCTAAGAAAAAGGACATG-3′

STn410 5′-GGTTGATAGTATCAACAACAGAGTC-3′

The underlined are accessory sequences for amplifying the 59-bp sequence flanked by the two Ace2 binding motifs. The *eng1-TATA*::*ura4*$^+$ strain used for the construction of the *eng1-TATA*Δ strain (see below) was transformed with the *eng1-motif*ΔΔ construct prepared with these primers.

***eng1-TATA*Δ construction**

STn407 5′-CCAAACAATATGTTGCCTGTAC-3′

STn408 5′-cagtgggatttgtagctaagcttCCAAATTTGGAAACTTGGACAG-3′

STn409 5′-gtttcgtcaatatcacaagcttGTATCAAGGTTCGCTTTTCTAATTAC-3′

STn410 5′-GGTTGATAGTATCAACAACAGAGTC-3′

STn411 5′-CTGTCCAAGTTTCCAAATTTGGGTATCAAGGTTCGCTTTTCTAATTAC-3′

The lowercase letters denote accessory sequences for amplifying the *ura4*$^+$ cassette. First, an *eng1-TATA*::*ura4*$^+$ strain was constructed with the former four primers, and the *ura4*$^+$ cassette was removed with the *eng1-TATA*Δ construct prepared with STn407, 408, 410, and 411. The underlined is complementary to the latter half of STn408 and was used to fuse the sequences flanking the putative *eng1* TATA box.

***eng1-ATG*Δ construction**

STn435 5′-CAATCAAATAACGCCAAAGTAATG-3′

| | |
|---|---|
| STn556 | 5′-cagtgggatttgtagctaagcttAGTTCCTAACAATAAAGTAAGTGAAAAG-3′ |
| STn557 | 5′-gtttcgtcaatatcacaagcttAGTTCCTATTTACGTTCTTTTATATTTGG-3′ |
| STn465 | 5′-AAGTAAAGCGAGCAGACATC-3′ |
| STn407 | 5′-CCAAACAATATGTTGCCTGTAC-3′ |
| STn421 | 5′-AGTTCCTAACAATAAAGTAAGTGAAAAG-3′ |
| STn544 | 5′-CTTTATTGTTAGGAACTAGTTCCTATTTACGTTCTTTTA-3′ |
| STn482 | 5′-AACAGCATCCTCAACATATCC-3′ |

The lowercase letters denote accessory sequences for amplifying the *ura4*$^{+}$ cassette. First, an *eng1-ATG*::*ura4*$^{+}$ strain was constructed with the former four primers, and the *ura4*$^{+}$ cassette was removed with the *eng1-ATG*Δ construct prepared with the latter four primers. The underlined is partially complementary to STn421 and was used to fuse the sequences flanking the first ATG of the *eng1* ORF.

***Peng1-ade1* construction**

| | |
|---|---|
| STn429 | 5′-ACGAAATTACGCACTCGAATG-3′ |
| STn430 | 5′-ggggatccgtcgacctgcagcgtacgaGCTTTCGTTACAGCGAGCAA-3′ |
| STn420 | 5′-gtttaaacgagctcgaattcatcgatATTTAACTACCTAGCTACTTAACGTC-3′ |
| STn421 | 5′-AGTTCCTAACAATAAAGTAAGTGAAAAG-3′ |
| STn431 | 5′-CTTTTCACTTACTTTATTGTTAGGAACTATGGAACCCATTATTGCGTTG-3′ |
| STn433 | 5′-CTTTCTAAAGCTTCAAAGGCTTC-3′ |

The lowercase letters denote accessory sequences for amplifying the *natMX6* cassette. The underlined is complementary to STn421 and was used to fuse the *Peng1* and *ade1* ORF fragments.

***Peng1-arg3* construction**

| | |
|---|---|
| STn423 | 5′-CAATGTATTGGTTCTTTAAAGAATGG-3′ |
| STn424 | 5′-ggggatccgtcgacctgcagcgtacgaTTTCAATTGCAAACTGAATAATTCTTC-3′ |
| STn420 | 5′-gtttaaacgagctcgaattcatcgatATTTAACTACCTAGCTACTTAACGTC-3′ |
| STn421 | 5′-AGTTCCTAACAATAAAGTAAGTGAAAAG-3′ |
| STn425 | 5′-CTTTTCACTTACTTTATTGTTAGGAACTATGTCTTTCAAAAAATTTCCTCGTC-3′ |
| STn427 | 5′-TGTAAAACGTTATTGGCATCAC-3′ |

The lowercase letters denote accessory sequences for amplifying the *natMX6* cassette. The underlined is complementary to STn421 and was used to fuse the *Peng1* and *arg3* ORF fragments.

***Pcfh4-ade1* construction**

| | |
|---|---|
| STn429 | 5′-ACGAAATTACGCACTCGAATG-3′ |
| STn430 | 5′-ggggatccgtcgacctgcagcgtacgaGCTTTCGTTACAGCGAGCAA-3′ |
| STn533 | 5′-gtttaaacgagctcgaattcatcgatATACACGAAAGATATTAAGGAAGATCTG-3′ |
| STn534 | 5′-CAACGCAATAATGGGTTCCATGGAAGTCACAGAGTCG-3′ |
| STn524 | 5′-ATGGAACCCATTATTGCGTTG-3′ |
| STn433 | 5′-CTTTCTAAAGCTTCAAAGGCTTC-3′ |

The lowercase letters denote accessory sequences for amplifying the *natMX6* cassette. The underlined is complementary to STn524 and was used to fuse the *Pcfh4* and *ade1* ORF fragments.

***Padg3-ade1* construction**

| | |
|---|---|
| STn429 | 5′-ACGAAATTACGCACTCGAATG-3′ |
| STn430 | 5′-ggggatccgtcgacctgcagcgtacgaGCTTTCGTTACAGCGAGCAA-3′ |
| STn535 | 5′-gtttaaacgagctcgaattcatcgatACATCGTTCATCTTCTGTAAAC-3′ |

STn536 5′-<u>CAACGCAATAATGGGTTCCAT</u>ACCCAAAAATAAAAAATTGTATTTATTGAAAG-3′

STn524 5′-ATGGAACCCATTATTGCGTTG-3′

STn433 5′-CTTTCTAAAGCTTCAAAGGCTTC-3′

The lowercase letters denote accessory sequences for amplifying the *natMX6* cassette. The underlined is complementary to STn524 and was used to fuse the *Padg3* and *ade1* ORF fragments.

***Pcdc22-ade1* construction**

STn429 5′-ACGAAATTACGCACTCGAATG-3′

STn430 5′-ggggatccgtcgacctgcagcgtacgaGCTTTCGTTACAGCGAGCAA-3′

STn527 5′-gtttaaacgagctcgaattcatcgatATTGACAACATCAAAAAGTAATATAAAGAAC-3′

STn528 5′-<u>CAACGCAATAATGGGTTCCAT</u>CGTGTTTGTTCTAGTTTGTAAG-3′

STn524 5′-ATGGAACCCATTATTGCGTTG-3′

STn433 5′-CTTTCTAAAGCTTCAAAGGCTTC-3′

The lowercase letters denote accessory sequences for amplifying the *natMX6* cassette. The underlined is complementary to STn524 and was used to fuse the *Pcdc22* and *ade1* ORF fragments.

***Pcdc18-ade1* construction**

STn429 5′-ACGAAATTACGCACTCGAATG-3′

STn430 5′-ggggatccgtcgacctgcagcgtacgaGCTTTCGTTACAGCGAGCAA-3′

STn531 5′-gtttaaacgagctcgaattcatcgatCTAGCTTAAAGTGTTCACTTGC-3′

STn532 5′-<u>CAACGCAATAATGGGTTCCAT</u>ATCGATACTTTATAGTAACCCCG-3′

STn524 5′-ATGGAACCCATTATTGCGTTG-3′

STn433 5′-CTTTCTAAAGCTTCAAAGGCTTC-3′

The lowercase letters denote accessory sequences for amplifying the *natMX6* cassette. The underlined is complementary to STn524 and was used to fuse the *Pcdc18* and *ade1* ORF fragments.

***Ptos4-ade1* construction**

| | |
|---|---|
| STn429 | 5′-ACGAAATTACGCACTCGAATG-3′ |
| STn430 | 5′-ggggatccgtcgacctgcagcgtacgaGCTTTCGTTACAGCGAGCAA-3′ |
| STn525 | 5′-gtttaaacgagctcgaattcatcgatTATCGCCAGGAACCTAAAC-3′ |
| STn526 | 5′-CAACGCAATAATGGGTTCCATACTGGCCAAAGGAGTA-3′ |
| STn524 | 5′-ATGGAACCCATTATTGCGTTG-3′ |
| STn433 | 5′-CTTTCTAAAGCTTCAAAGGCTTC-3′ |

The lowercase letters denote accessory sequences for amplifying the *natMX6* cassette. The underlined is complementary to STn524 and was used to fuse the *Ptos4* and *ade1* ORF fragments.

***Padh1-ade1* construction**

| | |
|---|---|
| STn429 | 5′-ACGAAATTACGCACTCGAATG-3′ |
| STn430 | 5′-ggggatccgtcgacctgcagcgtacgaGCTTTCGTTACAGCGAGCAA-3′ |
| STn422 | 5′-gtttaaacgagctcgaattcatcgatGCCCTACAACAACTAAGAAAATG-3′ |
| STn353 | 5′-AATTCTCTTGCTTAAAGAAAAGCG-3′ |
| STn432 | 5′-CGCTTTTCTTTAAGCAAGAGAATTATGGAACCCATTATTGCGTTG-3′ |
| STn433 | 5′-CTTTCTAAAGCTTCAAAGGCTTC-3′ |

The lowercase letters denote accessory sequences for amplifying the *natMX6* cassette. The underlined is complementary to STn353 and was used to fuse the *Padh1* and *ade1* ORF fragments.

***eng1*:*T*$_{CYC1}$(*1*) construction**

| | |
|---|---|
| STn733 | 5′-CCCTCACGAATCACCAAG-3′ |
| STn740 | 5′-gtaagcgtgacataactaattaTAGTAATTAGAAAAGCGAACCTTG-3′ |
| STn741 | 5′-ctcgaaggctttaatttgcTAACCTAAGTAAGACACAACCTC-3′ |
| STn736 | 5′-AGTAGCAATATCAACACATCTTG-3′ |

The lowercase letters denote accessory sequences for amplifying the *T*$_{CYC1}$ terminator element.

***eng1*:*T*$_{CYC1}$(*1086*) construction**

| | |
|---|---|
| STn733 | 5′-CCCTCACGAATCACCAAG-3′ |
| STn744 | 5′-gtaagcgtgacataactaattaCATTTGAGATGTGGATTCAGTC-3′ |
| STn745 | 5′-ctcgaaggctttaatttgcCTTGTTGGAAGCAACACTTTC-3′ |
| STn736 | 5′-AGTAGCAATATCAACACATCTTG-3′ |

The lowercase letters denote accessory sequences for amplifying the *T*$_{CYC1}$ terminator element.

***eng1*:*T*$_{CYC1}$(*2247*) construction**

| | |
|---|---|
| STn733 | 5′-CCCTCACGAATCACCAAG-3′ |
| STn746 | 5′-gtaagcgtgacataactaattaGGTAGTGTGAGTAATTTTGTTCATG-3′ |
| STn747 | 5′-ctcgaaggctttaatttgcTATTTTGGTACTAATATTGAGTACATCC-3′ |
| STn736 | 5′-AGTAGCAATATCAACACATCTTG-3′ |

The lowercase letters denote accessory sequences for amplifying the *T*$_{CYC1}$ terminator element.

***eng1*:*T*$_{CYC1}$(*3369*) construction**

| | |
|---|---|
| STn733 | 5′-CCCTCACGAATCACCAAG-3′ |
| STn748 | 5′-gtaagcgtgacataactaattaAGCAGCAACAAGTGCAC-3′ |
| STn749 | 5′-ctcgaaggctttaatttgcTAAGCATGACCAAAGTCCG-3′ |
| STn736 | 5′-AGTAGCAATATCAACACATCTTG-3′ |

The lowercase letters denote accessory sequences for amplifying the $T_{CYC1}$ terminator element.

***cdc10* tagging**

STn662 5′-CAAAACAGTCTATGATTGACTCAG-3′

STn663 5′-ggggatccgtcgacctgcagcgtacgaTGCTTGATGTTCTTTAACAACAC-3′

STn664 5′-gtttaaacgagctcgaattcatcgatTAATATTGCTTTTTGTGGTTTACCAC-3′

STn665 5′-AGCCACACTATAACTCCATG-3′

The lowercase letters denote accessory sequences for amplifying an epitope tag sequence on the pFA6a plasmid.

---

**Primers for qPCR**

---

***ade6* (Fig. 5a and Extended Data Fig. 4a-d, 5a, 6c, 7d)**

STn360 5′-GAAAGATGCTGCCGTCATTTTAG-3′

STn361 5′-GCTGCGGTACGAGCATAAGTAAC-3′

***eng1-Promoter* (Fig. 6d and Extended Data Fig. 4b)**

STn435 5′-CAATCAAATAACGCCAAAGTAATG-3′

STn436 5′-GCGGAAATCTGGATGTCTTA-3′

***eng1-ORF* (+126) (Fig. 5a)**

STn671 5′-CCTTAAGGATACCAAGGATACCAAA-3′

STn672 5′-AAATAACTGGCTGGGTTGATAGTA-3′

***eng1-ORF* (+838) (Fig. 5a, 6d and Extended Data Fig. 4a,c,d, 6c, 7d)**

STn481 5′-CTTGTTGGAAGCAACACTTTC-3′

STn482 5′-AACAGCATCCTCAACATATCC-3′

***eng1-ORF* (+1669) (Fig. 5a)**

STn646 5′-TACTTTCCAAAGCACCGAATG-3′

| | |
|---|---|
| STn647 | 5′-TTCGGACGTTGATTCTTCATC-3′ |

***eng1-ORF*** **(+2580) (Fig. 5a, 6d)**

| | |
|---|---|
| STn648 | 5′-TCGACCACTTCCTCCATTAC-3′ |
| STn649 | 5′-TTGACTGTGCGGTAGTAGTTG-3′ |

***eng1-3′ UTR*** **(Fig. 5a)**

| | |
|---|---|
| STn673 | 5′-CAGATGTACTATACTTATTGCTTGTATGT-3′ |
| STn674 | 5′-AGGGTTTCAAGTTGAGAGTAGTT-3′ |

***SPAC343.20-Promoter*** **(Extended Data Fig. 4b)**

| | |
|---|---|
| STn437 | 5′-ATGCAATTGATTCAAACATGAAAG-3′ |
| STn438 | 5′-GAGTGGCTGGTTGATGAG-3′ |

***SPAC343.20-ORF*** **(Extended Data Fig. 4a)**

| | |
|---|---|
| STn483 | 5′-CCGACTTATACGCACTACAATC-3′ |
| STn484 | 5′-TCGGCTACTAGGAAACCTTTA-3′ |

***ade1*** **(Extended Data Fig. 4c,d, 6c, 7d)**

| | |
|---|---|
| STn542 | 5′-AGCAGGTTGTGCTTTAGTAG-3′ |
| STn543 | 5′-CTCCAACTGAGGTTCCATTAG-3′ |

***arg3*** **(Extended Data Fig. 4c)**

| | |
|---|---|
| STn540 | 5′-CGTTACCGACACTTGGATTT-3′ |
| STn541 | 5′-GGCAAGCTTCATGATCTCTC-3′ |

***cnt2*** **(Fig. 6d)**

| | |
|---|---|
| STn131 | 5′-ACTAATAACGGAATAGATCAAACAAGCTC-3′ |
| STn132 | 5′-TATTTAACCAGCAAATTCATAGATTTGAC-3′ |

---

The *ade6* and *cnt2* primer sets for qPCR have been described previously[4].

### Supplementary Table 4 | Hi-C read summary.

| Sample name | Sequenced reads | Both side aligned | PCR bias removed | Excluding repeat-derived reads | MapQ>30 | Total Usable reads (Removed potential self-ligation and undigested reads) | inter-chromosome | intra-chromosome |
|---|---|---|---|---|---|---|---|---|
| cdc25-22_20min_Hi-C | 30,217,703 | 28,898,718 | 27,587,543 | 23,835,356 | 21,304,120 | 15,097,512 | 1,078,765 | 14,018,747 |
| cdc25-22_30min_Hi-C | 32,306,510 | 30,930,346 | 29,472,065 | 25,433,256 | 22,798,233 | 16,186,900 | 1,019,544 | 15,167,356 |
| cdc25-22_40min_Hi-C | 26,611,842 | 25,462,577 | 24,301,967 | 20,963,413 | 18,779,346 | 13,800,458 | 775,624 | 13,024,834 |
| cdc25-22_50min_Hi-C | 30,916,047 | 29,576,854 | 28,194,247 | 24,379,378 | 21,745,676 | 16,264,908 | 830,040 | 15,434,868 |
| cdc25-22_60min_Hi-C | 29,693,529 | 28,355,136 | 27,085,048 | 23,477,633 | 20,851,270 | 15,475,448 | 850,839 | 14,624,609 |
| cdc25-22_70min_Hi-C | 28,495,966 | 27,159,483 | 26,030,539 | 22,582,790 | 20,014,836 | 14,614,414 | 971,840 | 13,642,574 |
| cdc25-22_eng1-motif_20min_Hi-C | 29,152,932 | 27,721,973 | 21,236,419 | 17,828,962 | 14,957,560 | 10,169,719 | 600,852 | 9,568,867 |
| cdc25-22_eng1-motif_50min_Hi-C | 39,773,453 | 37,977,602 | 27,185,543 | 22,594,671 | 19,201,065 | 13,880,501 | 612,454 | 13,268,047 |
| cdc25-22_eng1-TATA_20min_Hi-C | 32,345,512 | 31,040,512 | 23,168,484 | 19,432,547 | 16,540,663 | 11,274,234 | 679,973 | 10,594,261 |
| cdc25-22_eng1-TATA_50min_Hi-C | 38,698,636 | 37,082,988 | 26,851,057 | 22,488,478 | 18,982,537 | 13,861,371 | 529,226 | 13,332,145 |
| cdc25-22_Peng1-ade1_20min_Hi-C | 22,416,841 | 21,404,046 | 18,304,175 | 15,578,038 | 13,053,775 | 8,779,266 | 430,462 | 8,348,804 |
| cdc25-22_Peng1-ade1_50min_Hi-C | 46,617,976 | 44,630,422 | 33,063,435 | 28,040,364 | 23,980,735 | 16,977,662 | 625,517 | 16,352,145 |
| cdc25-22_Peng1-arg3_20min_Hi-C | 31,933,318 | 30,233,190 | 22,485,923 | 18,856,399 | 15,760,284 | 10,855,299 | 617,873 | 10,237,426 |
| cdc25-22_Peng1-arg3_50min_Hi-C | 20,361,745 | 19,400,871 | 18,715,644 | 16,263,328 | 13,146,966 | 9,775,295 | 446,366 | 9,328,929 |
| cdc25-22_Peng1motif-ade1_20min_Hi-C | 20,318,883 | 19,191,174 | 16,594,216 | 14,059,093 | 12,026,410 | 8,308,077 | 483,803 | 7,824,274 |
| cdc25-22_Peng1motif-ade1_50min_Hi-C | 44,148,969 | 42,387,819 | 31,688,868 | 26,640,541 | 22,781,153 | 16,603,918 | 678,839 | 15,925,079 |
| cdc25-22_Peng1TATA-ade1_20min_Hi-C | 24,285,915 | 22,953,049 | 19,472,805 | 16,416,142 | 13,781,794 | 9,521,634 | 562,463 | 8,959,171 |
| cdc25-22_Peng1TATA-ade1_50min_Hi-C | 41,197,173 | 39,153,095 | 28,263,906 | 23,540,768 | 20,064,878 | 14,833,950 | 670,742 | 14,163,208 |
| cdc25-22_Pcfh4-ade1_20min_Hi-C | 32,464,988 | 30,966,373 | 22,815,444 | 19,355,037 | 16,291,016 | 11,138,514 | 627,000 | 10,511,514 |
| cdc25-22_Pcfh4-ade1_50min_Hi-C | 30,656,634 | 29,912,509 | 20,958,994 | 17,934,492 | 13,361,880 | 8,221,623 | 388,220 | 7,833,403 |
| cdc25-22_Padg3-ade1_20min_Hi-C | 29,973,898 | 28,718,025 | 21,392,416 | 17,933,193 | 15,111,137 | 10,241,778 | 607,887 | 9,633,891 |
| cdc25-22_Padg3-ade1_50min_Hi-C | 37,103,101 | 35,566,793 | 26,038,126 | 21,684,646 | 18,458,492 | 13,369,293 | 681,080 | 12,688,213 |
| cdc25-22_Pcdc22-ade1_20min_Hi-C | 26,376,431 | 25,233,579 | 18,829,367 | 15,873,092 | 13,530,327 | 8,976,946 | 527,016 | 8,449,930 |
| cdc25-22_Pcdc22-ade1_50min_Hi-C | 30,691,475 | 29,788,571 | 20,361,254 | 17,321,947 | 12,987,894 | 7,829,043 | 395,967 | 7,433,076 |
| cdc25-22_Pcdc18-ade1_20min_Hi-C | 30,912,043 | 29,654,048 | 22,076,006 | 18,420,518 | 15,618,305 | 10,552,366 | 660,995 | 9,891,371 |
| cdc25-22_Pcdc18-ade1_50min_Hi-C | 33,238,625 | 32,302,318 | 21,947,510 | 18,329,740 | 13,526,820 | 8,420,020 | 437,561 | 7,982,459 |
| cdc25-22_Ptos4-ade1_20min_Hi-C | 30,166,133 | 28,770,850 | 21,324,746 | 17,455,785 | 14,792,820 | 10,050,468 | 624,064 | 9,426,404 |
| cdc25-22_Ptos4-ade1_50min_Hi-C | 35,856,452 | 34,361,823 | 25,255,306 | 20,623,357 | 17,419,846 | 12,569,788 | 589,334 | 11,980,454 |
| cdc25-22_Padh1-ade1_20min_Hi-C | 19,594,359 | 18,508,658 | 16,040,877 | 13,610,798 | 11,454,458 | 7,860,321 | 514,843 | 7,345,478 |
| cdc25-22_Padh1-ade1_50min_Hi-C | 36,739,374 | 35,964,092 | 23,617,610 | 19,936,506 | 14,483,906 | 8,551,885 | 460,943 | 8,090,942 |
| cdc25-22_eng1:Tcyc1(1)_20min_Hi-C | 33,173,487 | 31,822,812 | 24,359,207 | 19,865,973 | 16,544,667 | 11,489,251 | 614,196 | 10,875,055 |
| cdc25-22_eng1:Tcyc1(1)_50min_Hi-C | 28,914,594 | 27,321,306 | 20,620,878 | 16,582,828 | 13,765,285 | 9,864,385 | 397,894 | 9,466,491 |
| cdc25-22_eng1:Tcyc1(1086)_20min_Hi-C | 32,299,317 | 30,874,564 | 23,278,320 | 19,497,051 | 16,553,796 | 11,420,357 | 596,036 | 10,824,321 |
| cdc25-22_eng1:Tcyc1(1086)_50min_Hi-C | 28,494,266 | 27,094,938 | 19,990,526 | 16,604,298 | 13,852,550 | 10,003,494 | 392,710 | 9,610,784 |
| cdc25-22_eng1:Tcyc1(2247)_20min_Hi-C | 34,448,881 | 33,051,341 | 24,295,801 | 20,001,966 | 16,807,442 | 11,638,495 | 602,823 | 11,035,672 |
| cdc25-22_eng1:Tcyc1(2247)_50min_Hi-C | 33,235,788 | 31,909,646 | 23,780,046 | 19,471,328 | 16,283,353 | 11,657,066 | 455,843 | 11,201,223 |
| cdc25-22_eng1:Tcyc1(3369)_20min_Hi-C | 32,217,328 | 30,692,667 | 23,172,027 | 18,938,206 | 16,010,999 | 11,161,133 | 605,632 | 10,555,501 |
| cdc25-22_eng1:Tcyc1(3369)_50min_Hi-C | 35,176,529 | 33,619,774 | 24,987,559 | 20,352,449 | 17,233,030 | 12,525,205 | 497,791 | 12,027,414 |
| pFA_MboI_Hi-C | 13,392,907 | 12,821,630 | 12,577,647 | 10,977,861 | 9,794,624 | 8,133,979 | 1,196,848 | 6,937,131 |
| pFA-DSG_MboI_Hi-C | 13,625,254 | 12,984,395 | 12,733,890 | 11,402,492 | 10,139,230 | 8,256,959 | 1,081,288 | 7,175,671 |
| pFA_MboI-HinfI_Hi-C | 14,693,754 | 13,917,048 | 13,653,997 | 11,298,532 | 10,088,536 | 8,240,401 | 1,201,744 | 7,038,657 |

| | | | | | | | | |
|---|---|---|---|---|---|---|---|---|
| pFA-DSG_MboI-HinfI_Hi-C | 12,864,342 | 12,234,804 | 12,014,312 | 10,513,048 | 9,356,797 | 7,579,014 | 953,005 | 6,626,009 |
| pFA_MboI-HinfI-MluCI_Hi-C | 19,752,949 | 18,882,558 | 18,298,474 | 15,847,213 | 14,336,312 | 10,892,903 | 1,669,800 | 9,223,103 |
| pFA-DSG_MboI-HinfI-MluCI_Hi-C | 20,709,819 | 19,696,733 | 19,258,831 | 16,963,476 | 15,000,426 | 11,392,340 | 1,316,480 | 10,075,860 |
| wt_pFA_MboI_Hi-C | 15,733,401 | 15,730,634 | 15,324,291 | 12,999,888 | 11,954,613 | 9,538,153 | 1,267,593 | 8,270,560 |
| cut14-208_pFA_MboI_Hi-C | 16,865,576 | 16,001,745 | 15,577,364 | 13,535,514 | 12,546,638 | 9,846,380 | 1,579,240 | 8,267,140 |
| rad21-K1_pFA_MboI_Hi-C | 16,032,020 | 16,028,330 | 15,651,917 | 13,057,123 | 11,974,519 | 9,764,285 | 2,749,425 | 7,014,860 |
| wt_pFA-DSG_MboI-HinfI-MluCI_Hi-C | 22,573,941 | 21,667,695 | 21,064,952 | 18,449,584 | 16,935,684 | 12,517,110 | 1,347,988 | 11,169,122 |
| cut14-208_pFA-DSG_MboI-HinfI-MluCI_Hi-C | 13,458,971 | 12,881,667 | 12,535,424 | 11,034,859 | 10,159,147 | 7,214,701 | 934,363 | 6,280,338 |
| rad21-K1_pFA-DSG_MboI-HinfI-MluCI_Hi-C | 15,870,246 | 15,867,264 | 15,444,436 | 13,022,695 | 11,916,481 | 8,959,959 | 2,176,885 | 6,783,074 |
| wt_pFA-DSG_MboI-HinfI-MluCI_Hi-C_NovaSeq | 201,536,272 | 194,121,149 | 156,018,991 | 134,163,159 | 126,837,179 | 94,005,272 | 10,180,120 | 83,825,152 |
| cut14-208_pFA-DSG_MboI-HinfI-MluCI_Hi-C_NovaSeq | 123,956,220 | 118,930,842 | 96,873,588 | 83,836,647 | 79,408,286 | 56,647,189 | 7,330,472 | 49,316,717 |

Read numbers remaining after the respective filtering processes are summarized as described previously[5].
